# In silico prediction of structure and function for a large family of transmembrane proteins that includes human Tmem41b

**DOI:** 10.1101/2020.06.27.174763

**Authors:** Shahram Mesdaghi, David L. Murphy, Filomeno Sánchez Rodríguez, J. Javier Burgos-Mármol, Daniel J. Rigden

## Abstract

Recent strides in computational structural biology have opened up an opportunity to understand previously mysterious uncharacterised proteins. The under-representation of transmembrane proteins in the Protein Data Bank highlights the need to apply new and advanced bioinformatics methods to shed light on their structure and function. This study focuses on such a family; transmembrane proteins containing the Pfam domain PF09335 (‘SNARE_ASSOC’/‘VTT ‘/‘Tvp38’). One prominent member, Tmem41b, has been shown to be involved in early stages of autophagosome formation and is vital in mouse embryonic development. Here we use evolutionary covariance-derived information not only to construct and validate *ab initio* models but also to make domain boundary predictions and infer local structural features. The results from the structural bioinformatics analysis of Tmem41b and its homologues show that they contain a tandem repeat that is clearly visible in evolutionary covariance data but much less so by sequence analysis. Furthermore, cross-referencing of other prediction data with the covariance analysis shows that the internal repeat features 2-fold rotational symmetry. *Ab initio* modelling of Tmem41b reinforces these structural predictions. Local structural features predicted to be present in Tmem41b are also present in Cl^−^/H^+^ antiporters. These results together strongly point to Tmem41b and its homologues as being transporters for an as-yet uncharacterised substrate and possibly using H^+^ antiporter activity as its mechanism for transport.

## Introduction

Protein structural information is crucial for an understanding of protein function and evolution. Currently, there is only experimental structural data for a tiny fraction of the protein universe ^1^. Membrane proteins are encoded by 30% of the human genome ^2^ but they have only a 2% representation in the Protein Data Bank (PDB) ^3^. Membrane protein families are particularly poorly understood due to their associated experimental difficulties, such as the challenge of over-expression which can result in toxicity to host cells ^4^ as well as finding a suitable membrane mimetic to reconstitute the protein. Additionally, membrane proteins are much less conserved across species compared to water-soluble proteins ^5^ making sequence-based homologue identification a challenge, and in turn rendering homology modelling of these proteins more difficult. Membrane proteins can be grouped according to their interaction with various cell membranes; integral membrane proteins (IMP) are permanently anchored whereas peripheral membrane proteins transiently associate with cell membranes. IMPs that span the membrane are known as transmembrane proteins (TMEMs) as opposed to IMPs that are monotopic, associating with one side of the membrane ^6^.

The transmembrane protein Tmem41 family has two members, Tmem41a and Tmem41b, both sharing the PF09335 (‘SNARE_ASSOC’/ ‘VTT ‘/’Tvp38’) Pfam ^7^ domain. The profile of Tmem41b has recently risen due to experimental evidence pointing to it, being involved in macroautophagy regulation (making it a possible Atg i.e. an autophagy related protein) and lipid mobilisation ^8^. Other studies identify Tmem41b to be involved in motor circuit function with TMEM41B knockout *Drosophila* showing neuromuscular junction defects and aberrant motor neuron development in knockout Zebra fish ^9^. Also, it has been shown that in TMEM41B knockout HeLa cells there is an inhibition of Zika virus replication ^10^. Additionally, Tmem41b has been shown to be essential for mouse embryonic development: homozygous knockout mice embryos suffer early termination of their development after 7-8 weeks ^11^. Tmem41b is a structurally uncharacterised 291-residue protein found in the endoplasmic reticulum (ER) localising at the mitochondria-associated ER membranes ^11^. Disruption of the PF09335 domain by various residue substitutions ^12^ or its removal ^13^ results in inhibition of autophagosome formation and impaired lipid mobilisation in human embryonic kidney (HEK) cells.

Tmem41b homologues are present in all domains of life ^14^. The Pfam PF09335 domain was first identified in the *Saccharomyces cerevisiae* protein Tvp38 ^15^, and the authors concluded that it associates with the tSNAREs in Tlg2-containing compartments, suggesting a role in membrane transport. Investigations into the bacterial and archaeal prevalence of these proteins showed that 90% of bacterial species and 70% of archaeal species encoded proteins with the PF09335 domain ^16^. Bacterial and archaeal PF09335-containing proteins are collectively known as the *Death Effector Domain A* (DedA) family. Detailed studies of the *Escherichia coli* DedA proteins have been carried out. There are eight *E. coli* representatives of the DedA family (YqjA, YghB, YabI, Yoh, EcdedA, YdjX, YdjZ, and YqaA) having overlapping functions ^14,16^ with Ydjx and Ydjz being the most closely related to human Tmem41b in terms of sequence similarity ^16^. Phenotypically DedA knock-out *E. coli* cells display increased temperature sensitivity, cell division defects, activation envelope stress pathways, compromised proton motive force, sensitivity to alkaline pH and increased antibiotic susceptibility ^16,17^. As *E. coli* expresses multiple DedA homologues the redundancy protects the cells from the phenotypical effects of single or multiple knock-outs as long as only one DedA is expressed ^17^. *Borrelia burgdorferi* contains only one DedA protein in its genome and knockout cells display the same phenotype as the *E. coli* knockout strains. Interestingly *E. coli* knockout cells can be rescued with the *B. burgdorferi* homologue that shows only 19% sequence identity with YdjA. Attempts to rescue *B. burgdorferi* DedA knockout cells with *E. coli* homologues resulted in more complex observations, with different homologues rescuing different phenotypes ^16^. The functions of DedA have also been studied in the opportunistic pathogen *Pseudomonas aeruginosa* where it was concluded that DedA proteins are required for its low antibiotic susceptibility. *P. aeruginosa* DedA is able to rescue *E.coli* DedA knockout cells ^18^.

Until the structure of poorly characterised protein families such as Pfam family PF09335 can be elucidated experimentally, *ab initio* protein modelling can be used to predict a fold allowing for structure-based function inferences ^19^. Such methods have made huge strides recently due to the availability of contact predictions ^20^. The combined use of metagenomic databases to generate large multiple sequence alignments (MSAs) and deep machine learning algorithms to detect evolutionary co-variance has increased the prediction accuracy of residue-residue contacts in protein tertiary structure significantly (Campaña et al. 2019; Greener et al. 2019; Kandathil et al. 2019). Prediction of residue-residue contacts relies on the fact that each pair of contacting residues covaries during the course of evolution. The process of co-variation occurs as the properties of the two residues complement each other in order to maintain structural integrity of that local region. Therefore, if one residue from the pair is replaced other must also change to compensate ^24^. The link between two residues can be then reliably detected in MSAs using direct coupling analysis ^25^. There is a positive correlation between the number of correctly predicted contacts and the number of effective sequences (Neff) of an MSA, i.e. the more numerous and diverse sequences the more likely accurate contact predictions can be made ^26^. Recent machine learning algorithms use multi-layered neural networks and utilise the contact predictions from multiple evolutionary coupling contact prediction methods along with sequence profiles to build a co-variance or precision matrix which is used to predict the contacts of a given protein ^22,27^. The predicted contacts can be used for a range of analyses such as identification of domain boundaries ^28,29^, but their main application is contact-based modelling methods which can address larger targets than conventional fragment-assembly based *ab initio* methods ^30^.

Here we utilise state of the art methods to make structural predictions for Tmem41b from data derived from sequence, evolutionary covariance and *ab initio* modelling. We are able to demonstrate that Tmem41b and its homologues contain re-entrant loops (stretches of protein that enter the bilayer but exit on the same side of the membrane) as well as a pseudo-inverted repeat topology. The presence of both of these structural features strongly suggests that Tmem41b and its homologues are secondary active transporters for an uncharacterised substrate.

## Materials and Methods

### Pfam Database Screening

Searches were made against the Pfam-A_v32.0 ^7^ database using the HHPred server ^31^ with default parameters and eight iterations for MSA generation in the HHblits ^32^ stage.

### Contact Map Predictions

The MSAs were generated using Jackhmmer v3.3 ^33^ default parameters with five iterations against a custom metagenomic-enriched database. This database was a concatenation of: Uniref100 ^34^, EupathDB ^35^, the Marine Eukaryotic Reference Catalogue (MERC) ^36^, the Soil Reference Catalogue (SRC) (Steinegger et al., 2018), MGnify ^37^. DeepMetapsicov was used to generate contact predictions with ConKit (Simkovic, Thomas, and Rigden 2017) being utilised to visualise the contact maps. Custom modifications were introduced into ConKit in order to overlay additional prediction data (Mesdaghi and Sánchez Rodríguez, unpublished work).

### Other Prediction Data

Transmembrane helical topology predictions were obtained from the Topcons server ^39^ as well as the TMHMM server ^40^. Secondary structure predictions were made employing a local installation of PSIPRED v4.0 ^41^; disorder predictions were made utilising the IUPred2a server ^42^. Residue conservation scores were obtained from the ConSurf server ^43^. ConKit was also used to predict and visualise potential structural domain boundaries ^28,44^.

### Dataset for Custom Re-entrant Database

A non-redundant set of 56 re-entrant helix sequences was built by first retrieving all 714 TM proteins that contain at least one re-entrant loop from the PDBTM ^45^ and removing redundancy with a 40% identity threshold. The resulting 127 protein structures were split into their component chains, eliminating any chain lacking a re-entrant loop. The subsequent set of 188 unique re-entrant loop sequences were then filtered removing any sequences of less than ten residues and more than twenty, thereby ensuring the collection of sequences conformed to the length of typical re-entrant loops. The remaining 56 sequences were clustered, supplemented by candidate re-entrant sequences from the proteins studied here. Clustering was performed using CLANS ^46^ with the BLAST results (p-value cut-off threshold of 0.1) ^47^ used to calculate strengths of similarity.

### Model Building

*Ab initio* models were constructed using a local installation of DMPfold v1.0 ^22^ with default parameters. Conservation was mapped on to the models using the ConSurf server ^43^. Visualisation of models was achieved using PyMOL^48^. Angles between the N-terminal and C-terminal halves of the re-entrant helices were calculated using the software Interhlx with the interhelical angle reported being 180 degrees minus angle ϴ calculated by the program VGM ^49^. Visualisation of the orientation and the angle that separate the two helices utilised the *AngleBetweenHelices* PyMol plugin ^50^.

*Ab initio* models were queried against the PDBTM ^45^ using Dali v4.0 ^51^.

## Results & Discussion

### Sequence comparisons suggest Pfam families PF09335 and PF06695 are related

HHpred (Zimmermann et al., 2018) was used to screen Tmem41b against the Pfam database (El-Gebali et al., 2019) producing hits in the same region against both PF09335 and the Pfam domain PF06695 (‘Sm_multidrug_ex’). The hits are unambiguously indicative of homology: a probability of 99.4% with an E-value of 9E-17 for the PF09335 hit while for PF06695 the values are 98.3% and 2E-10 respectively. A HHpred search against the Pfam database using Tmem41b homologues, including the short archaeal sequence Mt2055 (UniProt code W9DY28) ^35^ returned similar results (Table 1). The Mt2055 sequence originates from the unpublished draft genome of the archaebacterium *Methanolobus tindarius DSM 2278.* For many of the subsequent analyses, the shorter archaeal sequence was used initially but the clear homology among this set of proteins means that inferences can be drawn across the group.

**Table 1.**
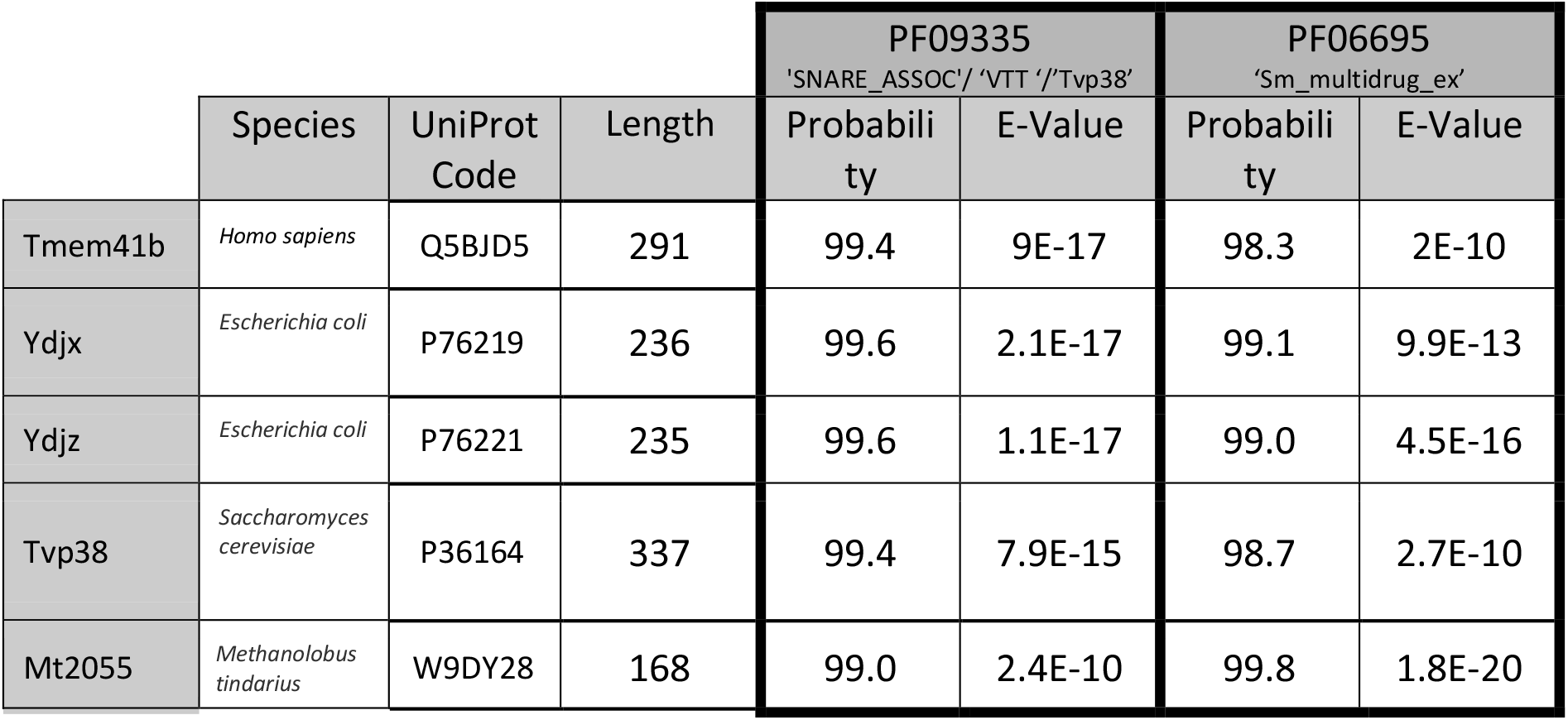
HHpred results for Tmem41b and homologues demonstrate homology between Pfam families PF09335 and PF06695.

### Sequence & contact prediction map analysis indicate that PF06695 is made up of a tandem repeat

Analysis of the HHPred results obtained for the archaeal protein Mt2055 revealed the presence of additional hits for both PF06695 and PF09335 Pfam domains, in which the C-terminal half of the domains aligned with the N-terminal half of the Archaea protein. For example, residues 1-69 of the archaeal protein aligned with residues 52-117 of the Pfam PF09335 profile with a probability of 74.15%. Interestingly, contact density analysis ^28,52^ supported the existence of a domain boundary around residue 60, in broad agreement with the HHpred results (see Fig. 1a). Furthermore, when the Mt2055 sequence was split at residue 60-61, the resulting N-terminal region of 60 residues and the C-terminal section of 79 residues could be aligned using HHalign ^53^ with a 78% probability and an E-value of 1.9E-3 (see Fig. 1b). Finally, examination of the map of predicted contacts for Mt2055 reveals features that are present in both the N- and C-terminal halves of the protein (see Fig. 1c). Taken together, these data strongly support the existence of a tandem repeat within the Mt2055 protein and hence across the PF06695 and PF09335 protein families.

**Fig 1.**
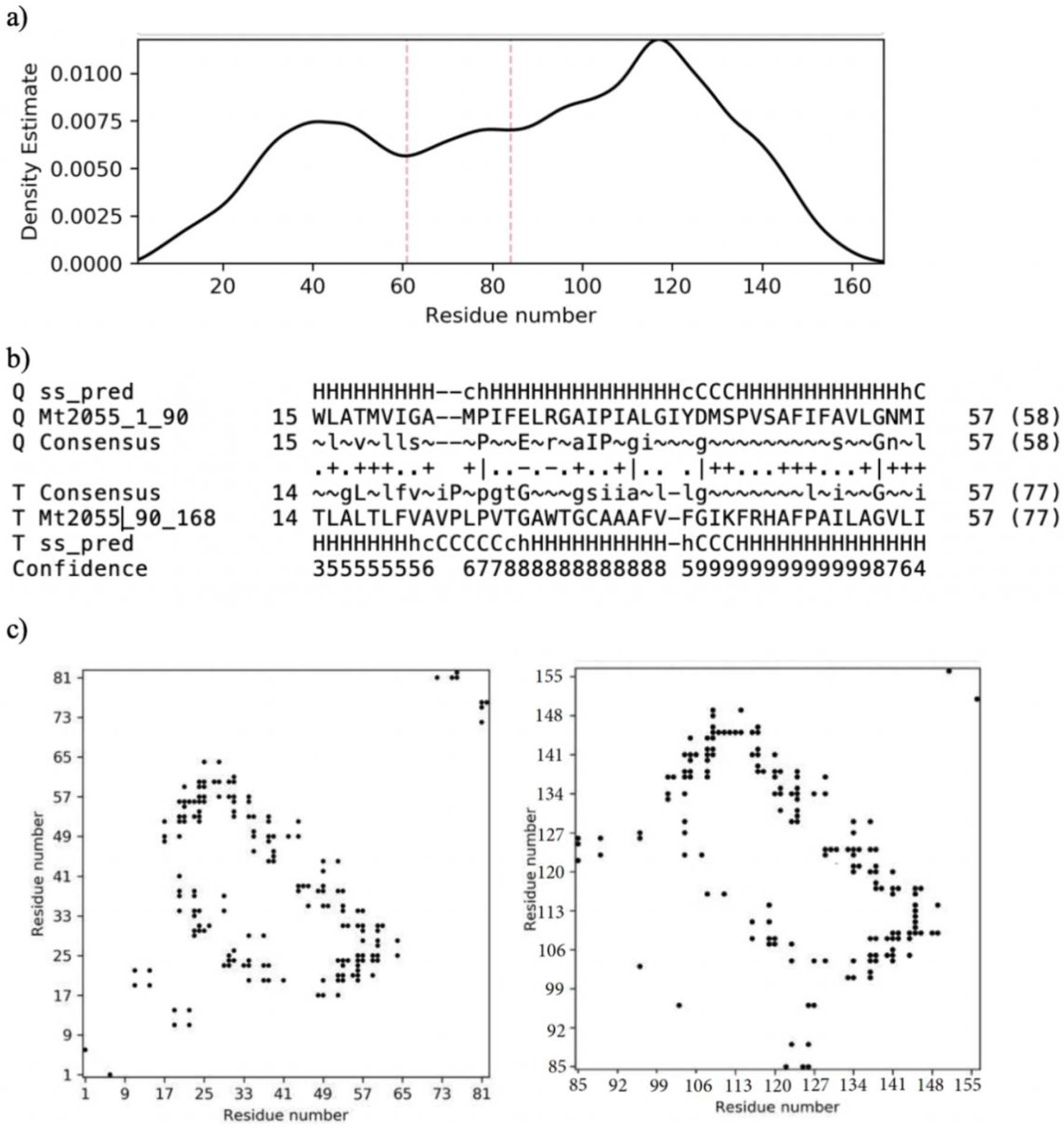
Mt2055 domain analysis. a) Contact density profile constructed by ConKit (Simkovic, Thomas, & Rigden, 2017) utilising DeepMetaPSICOV contact prediction. Solid black line represents contact density and dotted red lines mark density minima corresponding to possible domain boundaries. b) HHalign alignments for the N-terminal and C-terminal Mt2055 halves. c) Maps of predicted contacts generated by DeepMetaPSICOV and plotted using ConKit; left is N-terminal half (residues 1-84) and right is C-terminal half (residues 85-168). Black points represent predicted intramolecular contacts.

Interestingly, an equivalent sequence analysis with HHpred of Tmem41b or any of the other tested PF09335 homologues does not reveal a repeat. However, inspection of their corresponding predicted contact maps does reveal features repeated when N- and C-halves of the protein are compared (Fig. 2). Apparently, evolutionary divergence has removed all trace of the repeat sequence signal in bacterial and eukaryotic proteins, although the feature remains visible by evolutionary covariance analysis.

**Fig 2.**
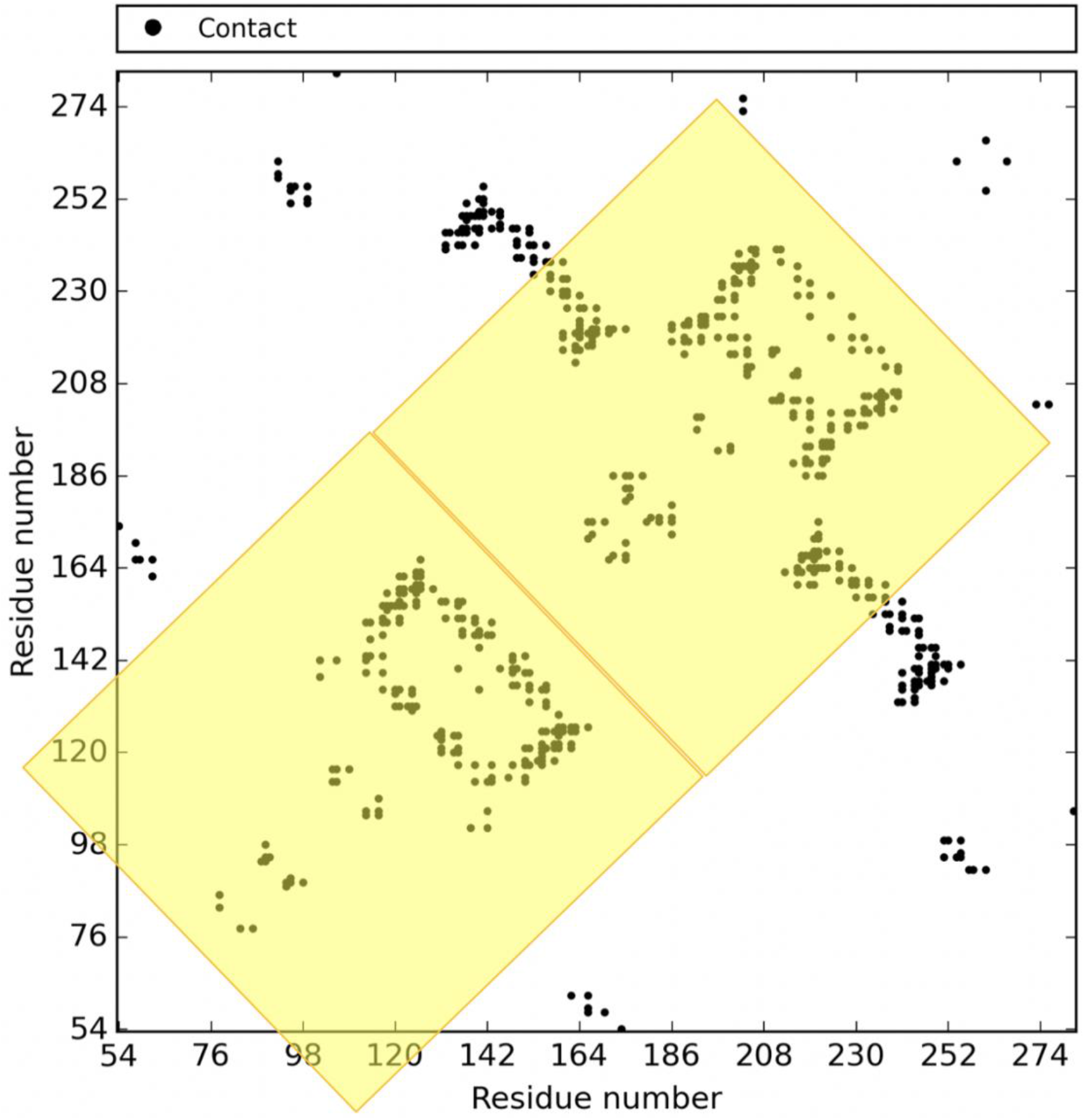
Tmem41b Contact map constructed using DeepMetaPSICOV (Kandathil et al., 2019) and plotted using Conkit (Simkovic, Thomas, et al., 2017). The highlighted areas represent repeat units that have been revealed through evolutionary covariance analysis.

### Study of contact map features, secondary structure predictions & membrane predictions show that PF09335 & PF06695 contain re-entrant loops and that the topology is made up of an inverted repeat

In order to assess the composition of the repeat identified, transmembrane helical topology predictions were carried out but gave inconsistent results for most proteins: only for the archaeal protein Mt2055 did all methods agree that four transmembrane helices were predicted to be present in the whole protein, two in each of the repeats (Table 2).

**Table 2.**
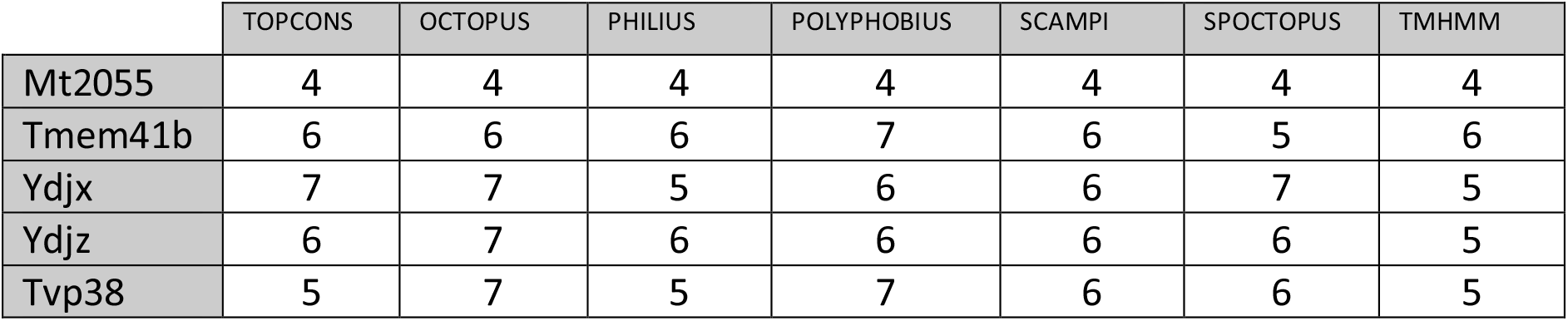
Predicted number of TM regions for PF09335/PF06695 homologues

To complicate matters further, secondary structure and contact predictions are incompatible with the TM helical predictions (Fig. 3). Analysis of the Mt2055 contact, membrane & secondary structure predictions showed that TM1 is split into two by the presence of a two-residue coil region halfway through the predicted TM helix. This might often indicate some kind of kink in the helix ^54^ but the explanation here is different. The contact predictions suggest that the N-terminal half of the helix makes contact with the C-terminal half. Structurally this could result in formation of a ‘V’ shaped helical structure or re-entrant loop embedded in the membrane. A similar pattern is repeated in TM3. The presence of re-entrant loops in a transmembrane protein strongly indicates a transporter or pore functionality since this structural feature has, hitherto, only been found in proteins of this kind ^55^.

**Fig 3.**
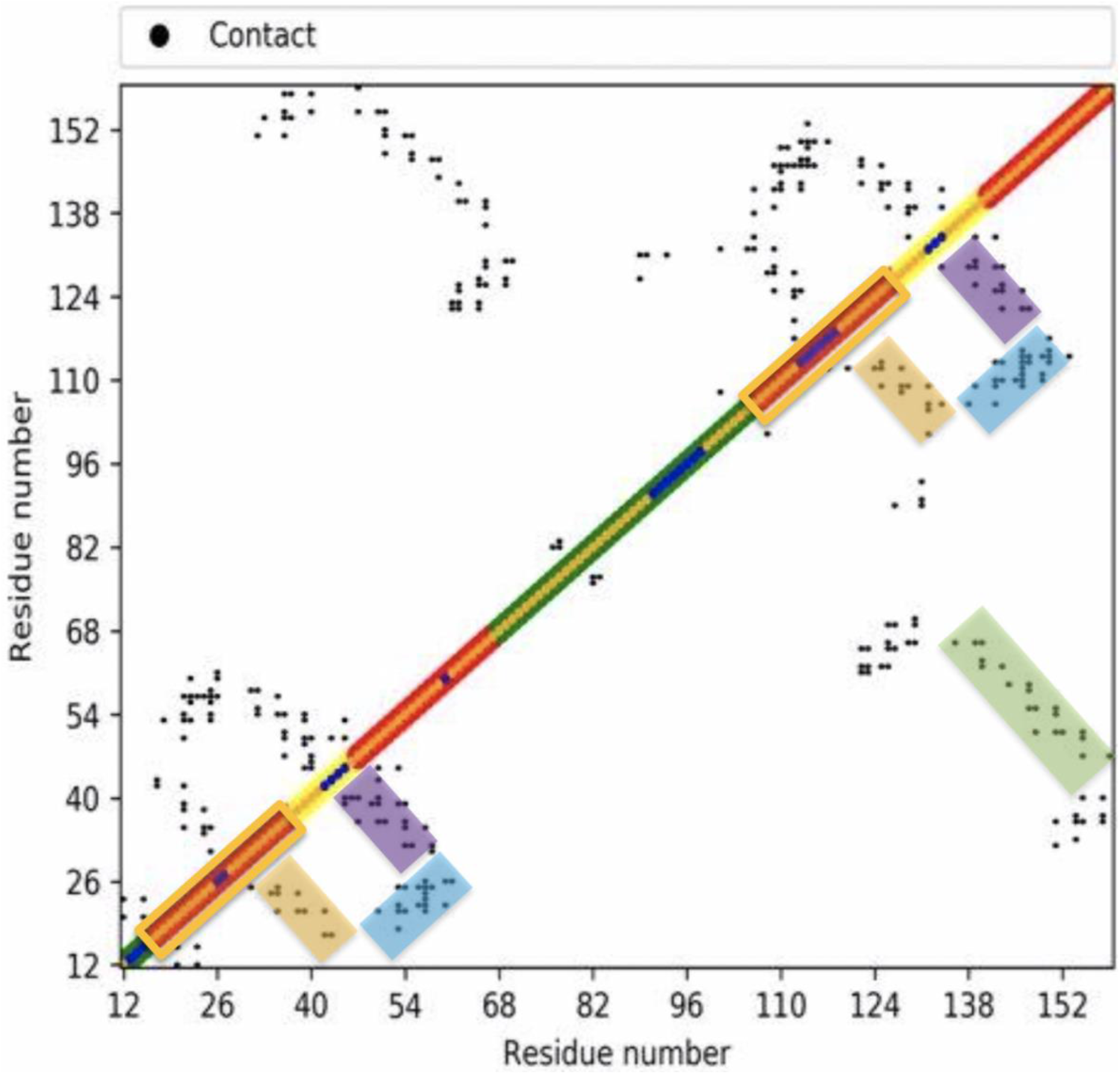
Mt2055 Contact map constructed using DeepMetaPSICOV predictions with TOPCONS (Tsirigos, Peters, Shu, Käll, & Elofsson, 2015) membrane prediction and PSIPRED (McGuffin, Bryson, & Jones, 2000) secondary structure predictions overlaid. The outer diagonals show the TOPCONS membrane prediction (red regions being predicted TM helices, green; inside cell, yellow; outside). The thin central diagonal is the secondary structure prediction (orange, helix; blue, coil). The figure was made with a modified (Mesdaghi and Sánchez Rodríguez, unpublished) version of ConKit. Orange boxes highlight proposed re-entrant loops i.e. regions predicted as TM 1 and TM 3 are in fact re-entrant loops by our analysis. Orange shaded regions highlight predicted contacts showing that N- and C-terminal halves of the re-entrant loop are in contact with one another. Green shading highlights contacts between predicted TM2 and TM4. Purple shading highlights contacts between N-terminal half of TM 2 and the C-terminal side of the re-entrant loop. Blue shading highlights contacts between the N-terminal half of TM 2 and N-terminal side of re-entrant loop.

The map also predicts that TM2 and TM4 transverse the membrane and are in contact with one another in three-dimensional space (green shading in figure 3), and, moreover, that half of each TM helix is intimately packed with a putative re-entrant loop (blue and purple shading in figure 3).

After the assignment of re-entrant loops and TM helices to the chain it became obvious that there was a helical region (to the N terminal of the second putative re-entrant loop) (residues 70-100) that required further exploration to determine its structural relevance. Amphipathic helices are sometimes present in TM proteins and are found lying on the membrane surface. This helix orientation results from an uneven distribution of polar and hydrophobic residues between the two faces of the helix ^56^. The amphipathic support vector machine prediction server, Amphipaseek ^57^ was used to predict regions of amphipathicity. Amphipaseek hinted at short amphipathic helices on the N-terminal side of each putative re-entrant loop (Fig. 4a). Performing the same prediction method on Tmem41b and its homologues does not suggest the presence of any amphipathic features although the corresponding regions in these homologues do show somewhat higher scores and are local maxima in the profiles (Fig. 4b, supplementary Figure 1).

**Fig 4.**
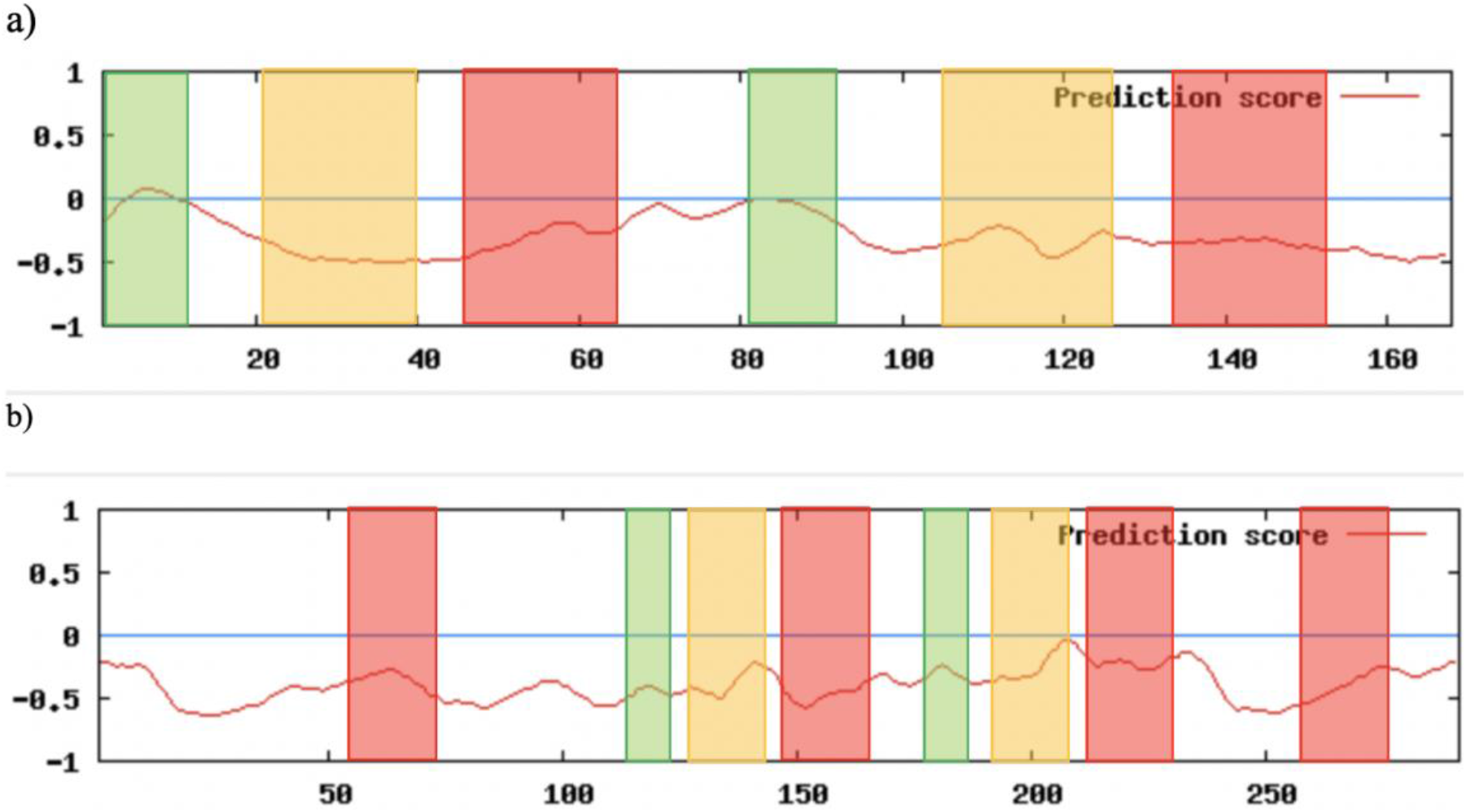
Amphipathic helix predictions for a) Mt2055 and b) Tmem41b. For both figures red shading indicates predicted TM helices, green shading highlights local maxima scores at the start of each repeat, and orange shading indicates putative position of re-entrant loops.

Using the predicted contact map data, a topology map can be generated (Fig. 5) that also satisfies secondary structure predictions as well as the results above that indicated the presence of an internal repeat. The constructed topology map reveals that the two units making up the tandem repeat show opposing membrane topology resulting in an inverse repeat structure ^58^.

**Fig 5.**
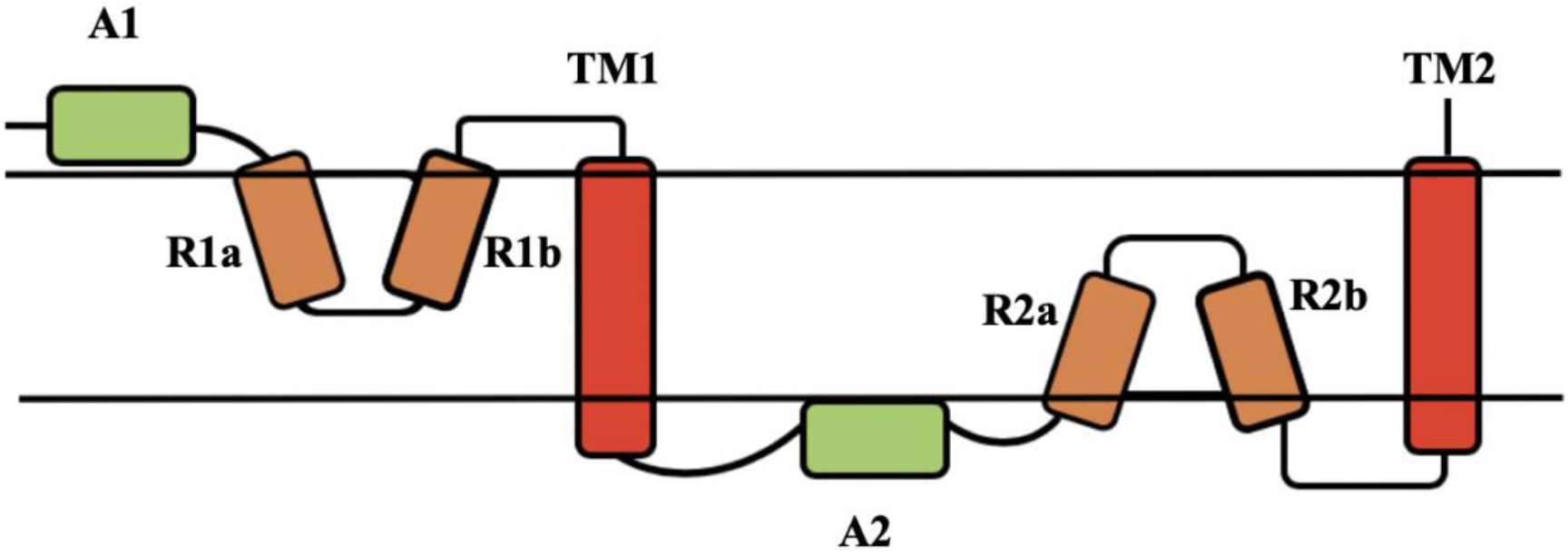
Predicted topology for Mt2055 based on contact, secondary structure, membrane & amphipathic predictions. Residue number increases from left to right. The protein domains are: A1 – amphipathic helix 1; R1a (N-terminal half of re-entrant helix 1; R1a (C-terminal half of re-entrant helix 1; TM1-transmembrane helix 1; A2 – amphipathic helix 2; R1a (N-terminal half of re-entrant helix 2; R2a (C-terminal half of re-entrant helix 2; TM2-transmembrane helix 2.

Predicted contact map features and secondary structure predictions for Tmem41b indicate that this protein also displays the same core topology of an inverted repeat made up of an amphipathic helix, a re-entrant loop and a TM helix. However, there are two additional TM helices on either side of the proposed conserved core topology (Fig. 6). Indeed, when examining the contact prediction maps for Tvp38, Ydjx and Ydjz, it can be concluded that they also have the inverted repeat topology shown in figure 5 but with additional TM helices as with Tmem41b.

**Fig 6.**
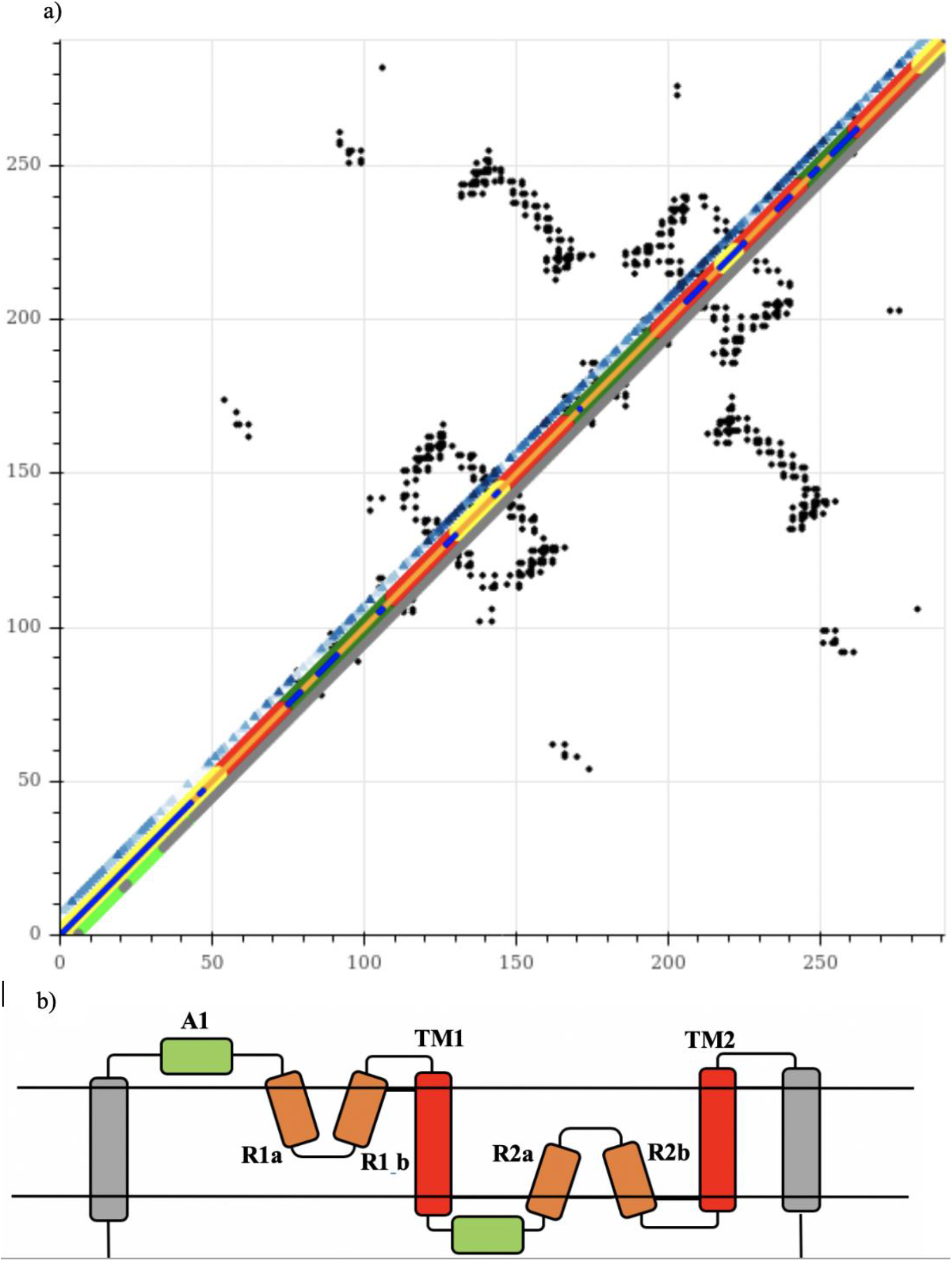
a) Tmem41b predicted contact map constructed using DeepMetaPSICOV with additional information overlaid on the diagonal. The outer diagonals show the TOPCONS membrane prediction (red regions being predicted TM helices, green; inside cell, yellow; outside). The thin central diagonal is the secondary structure prediction (orange, helix; blue, coil). Additionally, the upper diagonal contains a blue spectrum which indicates levels of sequence conservation from ConSurf (Ashkenazy et al., 2016) (the darker the blue the higher the level of conservation). Also, the grey (ordered) and lime green (disordered) lower diagonal utilises disorder predictions from IUPRED2a. b) Proposed topology for Tmem41b, with protein domains: A1 – amphipathic helix 1; R1a (N-terminal half of re-entrant helix 1); R1a (C-terminal half of re-entrant helix 1; TM1-transmembrane helix 1; A2 – amphipathic helix 2; R1a (N-terminal half of re-entrant helix 2); R2a (C-terminal half of re-entrant helix 2); TM2-transmembrane helix 2; with the presence of two-additional TM helices compared to Mt2055; Grey TM helices are additional helices to the core present in Tmem41b.

The sequence-structure relationships of re-entrant loops have been studied before ^55^ revealing that while TM helices have an even distribution of hydrophobic residues, re-entrant loops show an uneven distribution. Indeed, examination of the putative re-entrant loop sequences identified an inconsistent hydrophobicity distribution in both putative re-entrant loops of the homologues studied here (Fig. 7) with the C terminal side being more hydrophilic. Interestingly it is the residues of the C terminal side of the re-entrant loops that more conserved (Fig. 6a).

**Fig 7.**
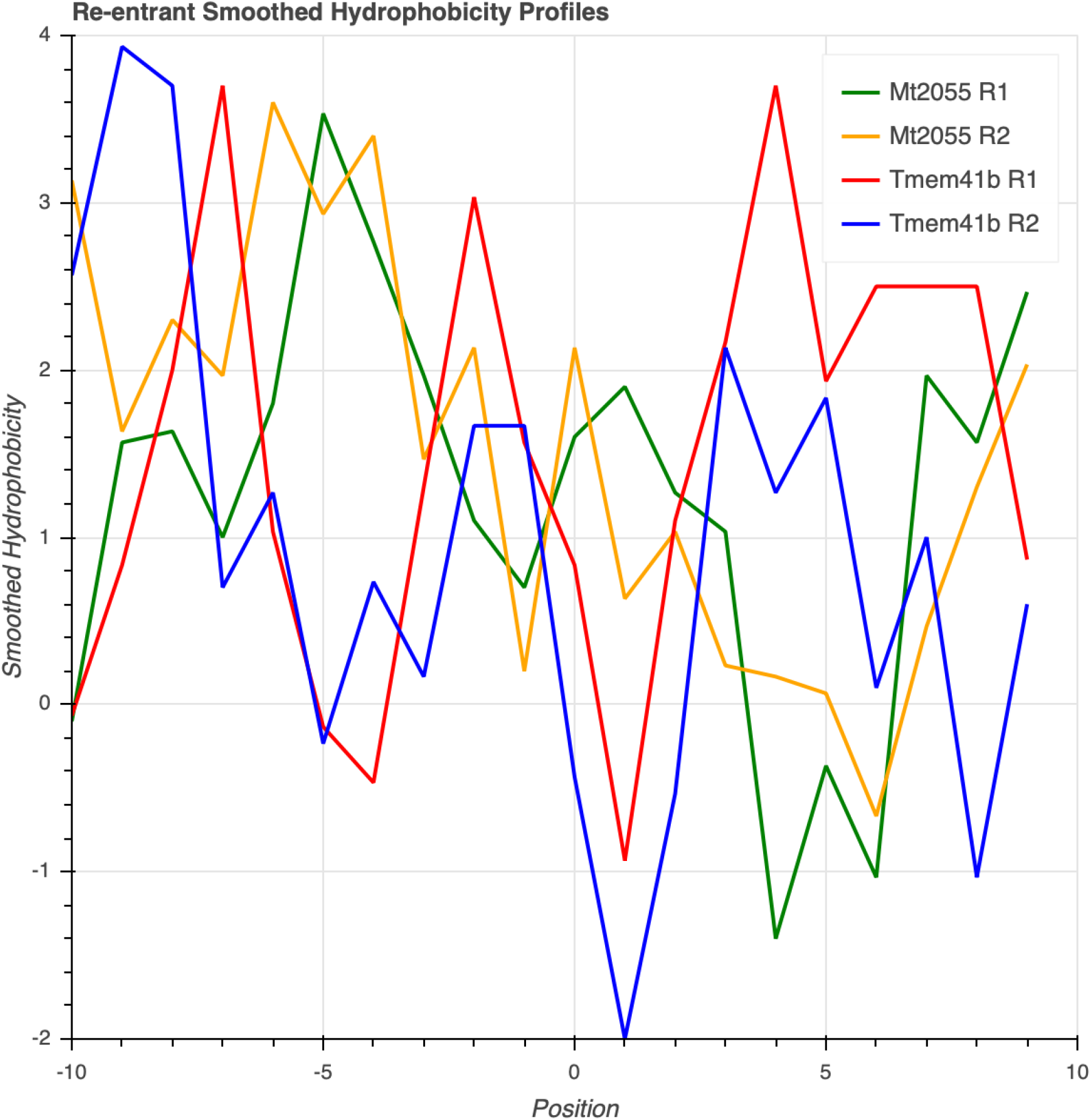
Smoothed hydrophobicity profiles for putative re-entrant loops of Mt2055 and Tmem41b. Smoothed the hydrophobicity distribution using a sliding window of three residues. For each position the mean hydrophobicity (Kyte & Doolittle, 1982) of the three positions covered by the window is calculated and assigned to the position at the center of the window. Positions are numbered by assigning the central residue (proline; see later) as 0.

### The predicted pseudo inverted repeat structure of Tmem41b and Mt2055 reinforces the hypothesis that they are transporters

It has been argued that the presence of symmetry in membrane proteins aids stability and provides the mechanics for conformational changes ^59^. Inverted pseudosymmetry topology of membrane proteins is only found in transporters and channels ^60^. The inverted symmetry is considered to allow access to functional residues or structures on both sides of the membrane and to provide stability for the proteins as symmetric polar ends of the TM helices that are able to face outside of the membrane ^61^. This is more favourable energetically as polar ends are not buried in the membrane. In chloride channels the antiparallel topology contributes to the selectivity of the hydrophobic anions as hydrophilic cations are not able to overcome the dielectric barrier of the membrane and require helix dipoles to be arranged in parallel ^62^.

The identification of inverse symmetry present in Mt2055 is exciting as it represents the smallest example of an integral membrane protein displaying this structural property. In order to possess the inverse repeat, each symmetrical unit must contain an odd number of TM helices in order to satisfy the gene duplication/fusion origin hypothesis for these proteins ^61^. Mt2055 only has one TM helix in each unit with the re-entrant loops and TM helices of each unit meeting within the membrane to contribute to the structurally conserved core of the putative transporter.

### Contact Map features assigned to re-entrant loops are also present in Cl^−^/H^+^ Antiporter predicted contact maps

In order to validate our interpretation of predicted contact map features as re-entrant helices, we examined experimentally characterised re-entrant loops in the PDBTM database^45^. A total of 56 non-redundant re-entrant helices were identified (see Methods). All 56 were clustered with the putative re-entrant loops from Mt2055 and the PF09335 homologues using relative E-values derived from an all-against-all BLAST run in CLANS ^46^ with a 0.1 p-value cut-off. The largest cluster contained 14 sequences, of which four were putative re-entrant sequences from the query proteins (Mt2055 R2, Ydjx R2, Ydjz R1 & Ydjz R2), seven (3org, 5tqq, 3nd0, 3det and 6coy) were re-entrant loop sequences from Cl^−^/H^+^ antiporters, one was from a boron exchanger (5l25), one from an electron transporter (2n4x) [albeit one classified as a member of the lysine exporter superfamily ^63^] and one from a mechanogated channel (5z10). Comparing the archaeal and Tmem41b contact map predictions with the maps derived from experimental data for all members of the largest cluster exposed the presence of similar features in the Cl^−^/H^+^ antiporters (Fig. 8). These features were also observed in predicted contact maps for the antiporters (Fig. 8a) supporting the inferences made for Mt2055 and Tmem41b. The matching features reflect the presence of a re-entrant loop and its packing against an adjacent transmembrane helix. The identification of these features in the predicted contact map strengthens the interpretation above made of the similar features in Tmem41b and relatives.

**Fig 8.**
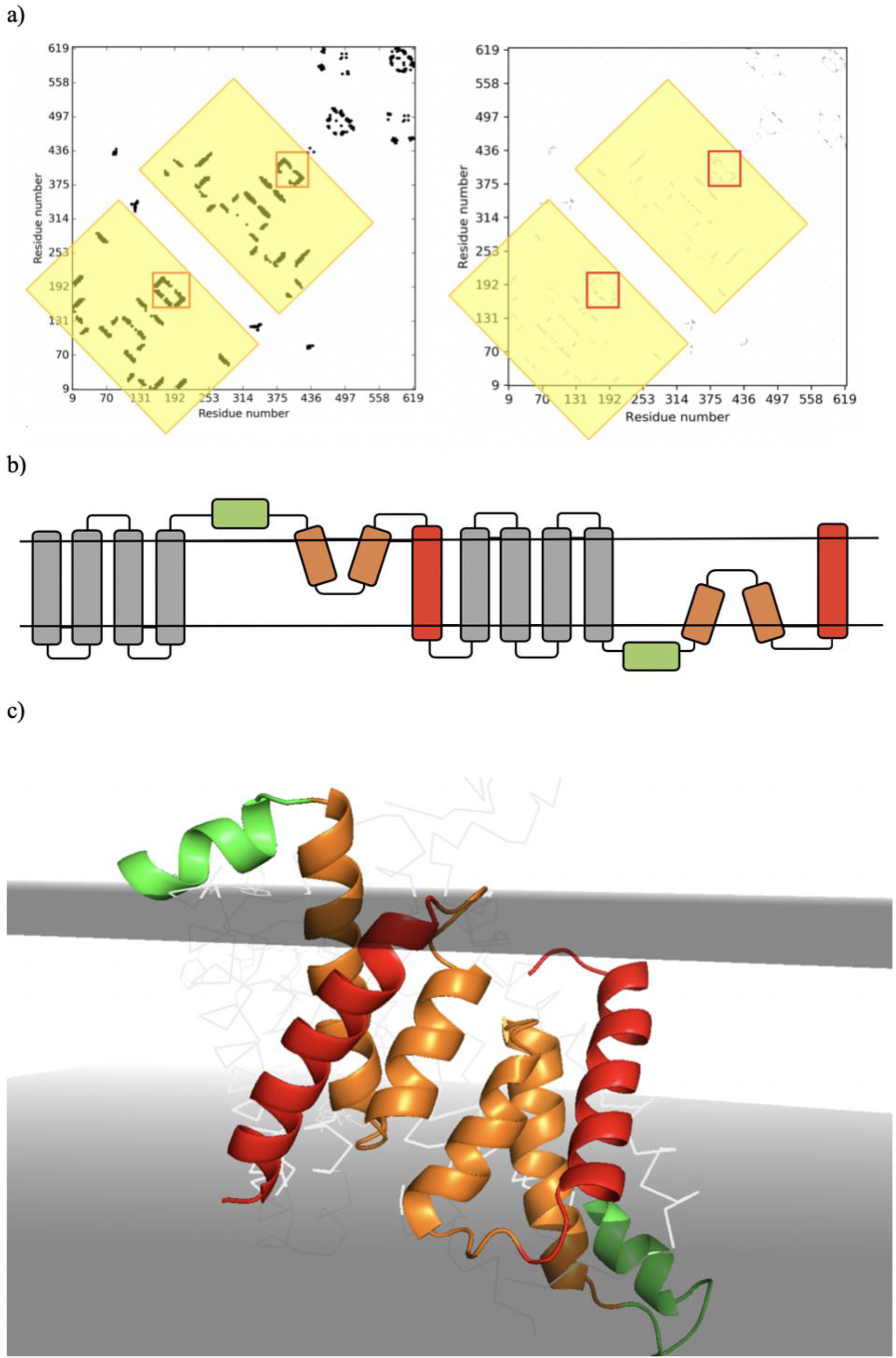
Structure information of the Cl^−^/H^+^ antiporter 3orgA a) Predicted Contact map with repeating units highlighted, contact map signature of re-entrant packed with TM helix in red boxes and corresponding regions in crystal structure shown in b); b) The Experimental Contact map obtained from the PDB structure with repeating units highlighted, key features in red boxes and corresponding regions in crystal structure shown d). c) Predicted topology; grey: TM Helices that are additional to the core; red: TM helices contributing to the formation of the core; orange; re-entrant loops contributing to the formation of the core; green: amphipathic helices contributing to the formation of the core. d) The 2-fold pseudo symmetry of the amphipathic/re-entrant loop/TM helix core inverted repeat structure of 3orgA with membrane positions obtained from PDBTM).

Additionally, the Cl^−^/H^+^ antiporter structures contain a similar inverted repeat as we propose for Tmem41b and homologues, resulting in pseudo-2-fold axis of symmetry running along the membrane ^61^. Again similarly, the Cl^−^/H^+^ antiporter 3orgA also contains the amphipathic helices on the N-terminal side of the re-entrant loops. The fact that the presence of the amphipathic helices is restricted only to 3orgA and that conservation analysis in the query proteins suggest that these features are not essential for function.

The similarities between the query proteins and the Cl^−^/H^+^ antiporters raise the possibility that the families studied here are, in fact, unsuspected distant homologues having this putative pore feature in common. In that regard it is relevant to recall a hypothesis that DedA proteins are H^+^ antiporters ^18^. However, the conserved and functionally important *proton* or *gating* glutamates that are responsible for H^+^ antiporter activity present in the Cl^−^/H^+^ antiporters ^64^ are absent in the proteins studied here. A recent study has identified key residues in the E. coli DedA protein Yqja that, when replaced in site directed mutagenesis experiments, it results in decreased proton motive force across the E. coli inner membrane^65^. Using the contact plot for Yqja, the essential residues (E39, D51, R130 and R136) map to the putative re-entrant loop regions. The N terminal side of the first re-entrant possesses E39 with the C terminal side possessing D51. R230 and R136 are similarly distributed on the second re-entrant loop (supplementary Figure 2).

### DMPfold modelling of the Archaea Mt2055 protein constructs a model that respects the predicted topology

Several authors have deposited structures of uncharacterised Pfam families in databases such as the Gremlin group ^26^ and the Jones group ^22^. The models retrieved for PF09335 and PF06695 are consistent but do not respect the predicted topology we have argued for. A quantitative comparison using TMalign ^66^ of Pfam PF09335 & PF06695 *ab initio* models constructed using DMPfold ^22^ pipeline shows a TM Score of 0.7 indicating that both models have the same fold (Fig. 9). Additionally, comparison of the *ab initio model* of PF06695 produced by the Gremlin group with either of the DMPfold models again results in a TM score of 0.7 (Fig. 9). This result reinforces the idea that PF09335 & PF06695 have the same fold.

**Fig 9.**
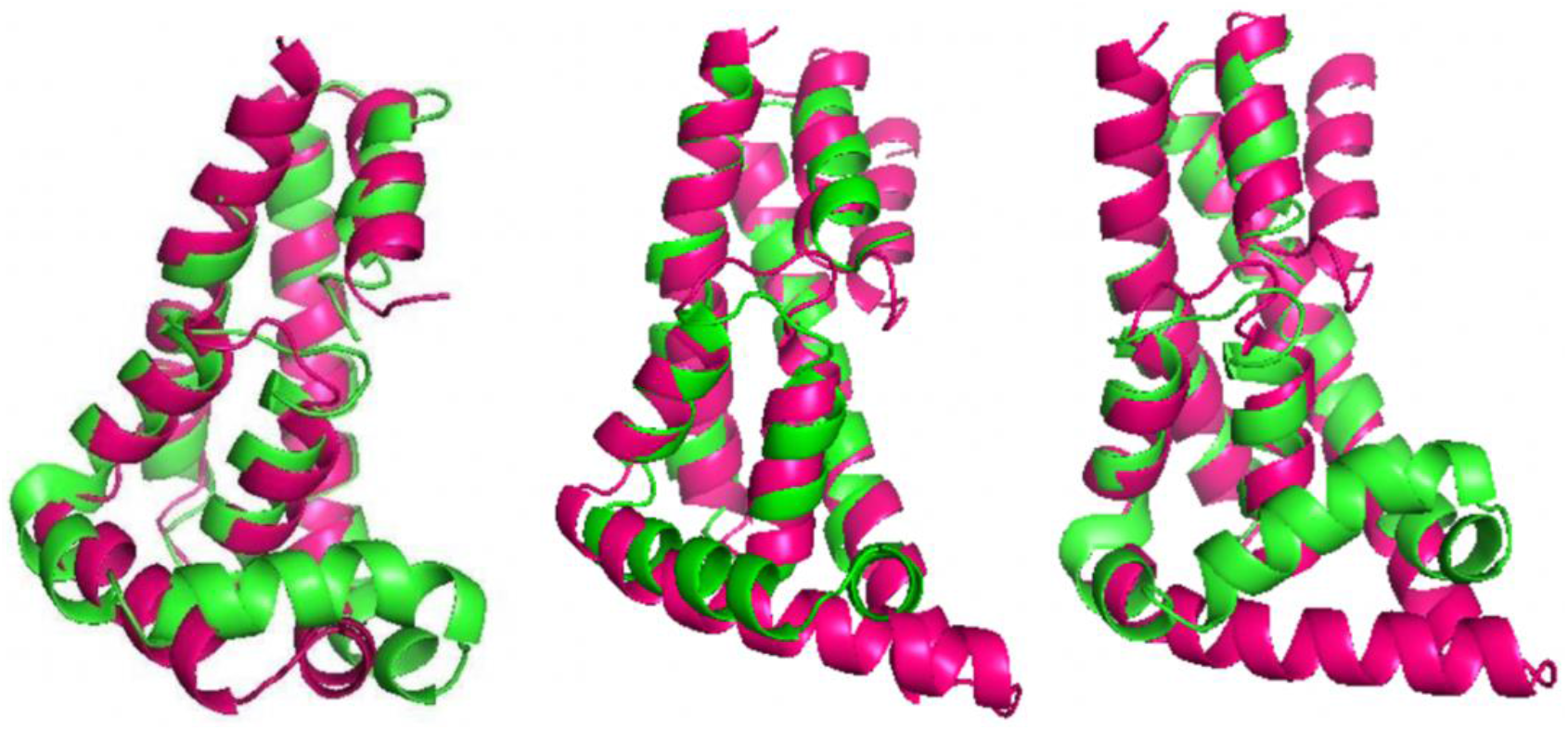
TMAlign superpositions of automated pipeline ab initio models made by Gremlin 26 and DMPFold 22. Left; DMPfold automated pipeline model for PF09335(pink) aligned with DMPfold model for PF06695(green) produces a TM score of 0.73 (normalised by the PF06695). Middle; Gremlin automated pipeline model for PF06995(pink) with the DMPfold PF09335 model (green) has a TM score of 0.75 (normalised by the DMPfold PF09335 model). Right; Gremlin automated pipeline model for PF06995(pink) with the DMPfold PF06695(green) has a TM score of 0.72 (normalised by the DMP PF06695 model).

However, it is important to note that the Pfam domain boundaries for PF09335/PF06695, which define the limits of previous modelling exercises, do not reflect a predicted conserved structural domain. The Pfam domain PF09335 covers the sequence from R1b to the halfway point of TM2 whereas PF06695 covers the sequence from R1b and the whole of TM2 (Supplementary Figure 3). The fact that the Pfam domain boundary does not cover the whole ‘functional unit’/core poses a problem for automated *ab initio* modelling pipelines attempting to model the PF09335 & PF06695 domains: the resulting models have half are-entrant loop on the N-terminal side and, in the case of PF09335, half a transmembrane helix as shown in figure 10.

**Fig 10.**
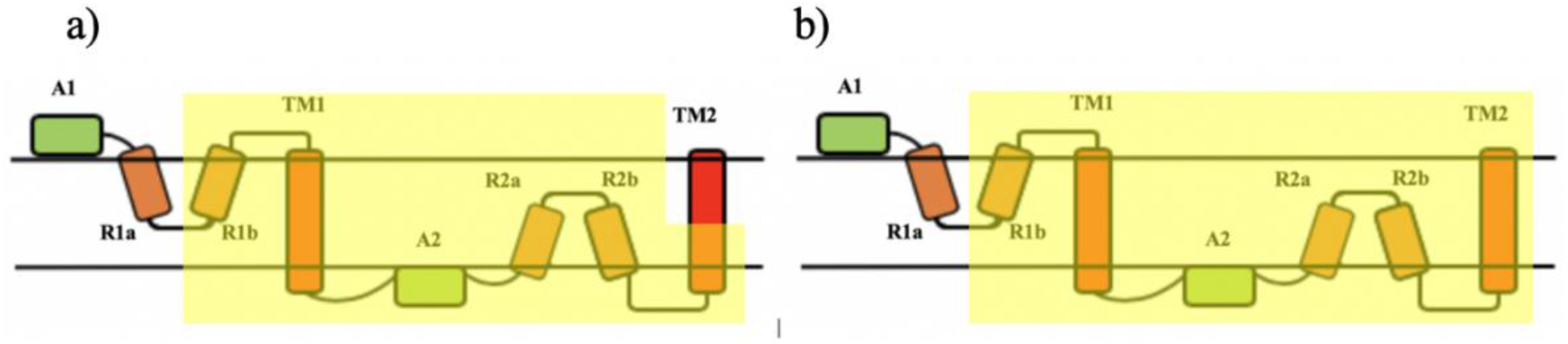
a) Proposed complete core topology for PF06695/PF09335 homologues with current PF09335 domain limits highlighted. b) Proposed complete core topology for PF06695/PF09335 homologues with current PF06695 domain limits highlighted.

Given the fact that available *ab initio* models were inconsistent with other data we constructed our own models of Mt2055 with DMPfold. The inclusion of metagenomic sequence data ^26,67,68^ was used to generate a large MSA lifting Neff value ^69^ from 1648 when using Uniref90 ^35^ to 7470 (Supplementary Figure 4).

The resulting output model (Fig. 11) contained all the predicted key features: two inversely symmetrical repeated units each possessing, in order, an amphipathic helix, a re-entrant loop and a TM helix. Evaluation of the model based on its satisfaction of predicted contacts ^29^, shows that 80% of the top *l* predicted contacts (where l is the length of the protein) are satisfied by the model contacts (Supplementary Figure 5). Additionally, when conservation is mapped on to the model using ConSurf ^43^ several stretches of conserved residues that are far apart in the sequence come together in three-dimensional space (Fig. 15). By comparison with the Cl^−^/H^+^ antiporter it can be hypothesised that the zones line a transmembrane pore and are responsible for substrate selectivity.

**Fig 11.**
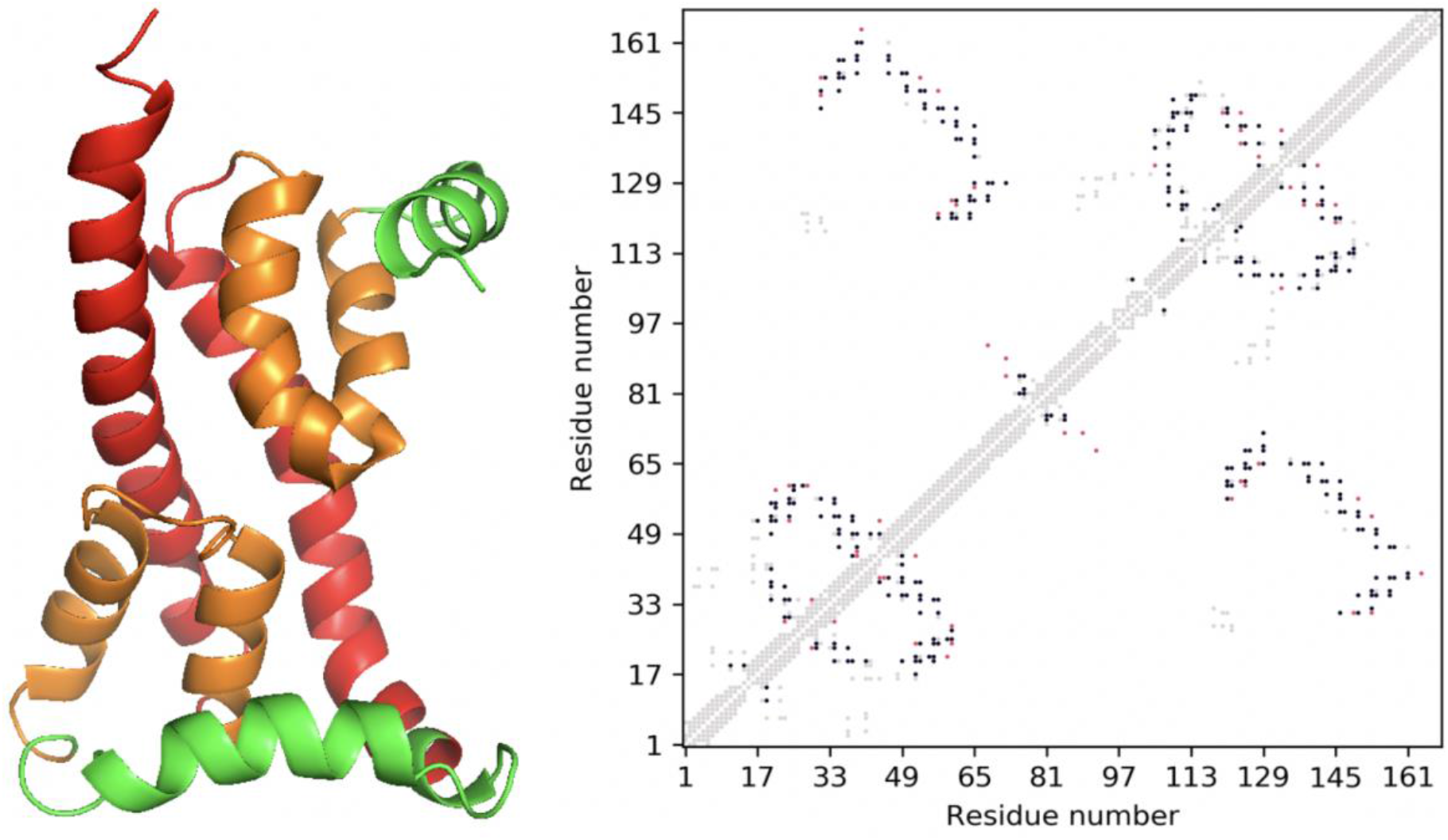
a) DMPfold model of Mt2055 (Green (amphipathic) red (TMHelix) and orange (re-entrant loop)) b) Comparison of model contact map with predicted contact map. Grey points are structural contacts of the model; black points are where the model and the contact predictions match; red points are predicted contacts not present in the model.

Dividing the model into two fragments according to the predicted domain boundary and then aligning the two portions using TMalign gave a TM-score of 0.66. This supports the proposal that the protein contains an inverted repeat structure with 2-fold rotational symmetry about the membrane (Fig. 12).

**Fig 12.**
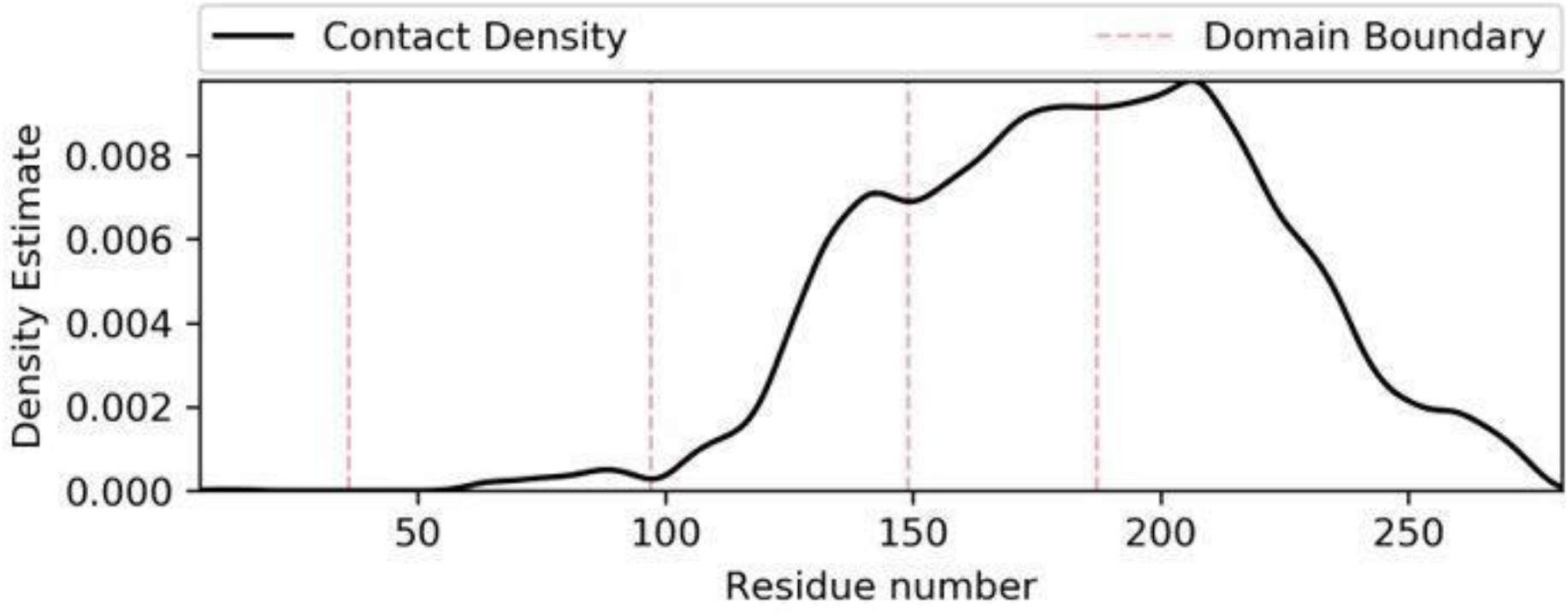
Predicted domain boundaries for Tmem41b based on predicted contact density. Disorder prediction data suggests that the first 40 residues are disordered. TM helix prediction data indicate the presence of a TM helix from positions 50-70 with the contact density profile showing that this region is not intimately packed with the rest of the protein.

### DMPfold modelling of the Tmem41b protein constructs a model that respects the predicted topology of the core region

Tmem41b, being significantly longer than Mt2055 and having additional secondary structure elements (Fig. 6), is a significantly more difficult modelling case. Indeed, modelling of the whole of the 291-residue Tmem41b proved unsuccessful due to large stretches lacking predicted secondary structure and the presence of a large number of contacts in the model that are not present in the contact predictions. However, the sequence could be truncated at the N-terminus by 100 residues since the first 40 residues were predicted to be intrinsically disordered (Fig. 6a) while the following 60 had few predicted contacts to the remainder of the protein: the predicted contact density profile (Supplementary Figure 6) indeed suggested the existence of a domain boundary at around residue 100. After removing 100 N-terminal residues, DMPfold produced a model in line with the predicted topology, having a good precision score (60% for top *l* contacts (Supplementary Figure 7)) and again showing clustering of conserved residues (Fig. 13).

**Fig 13.**
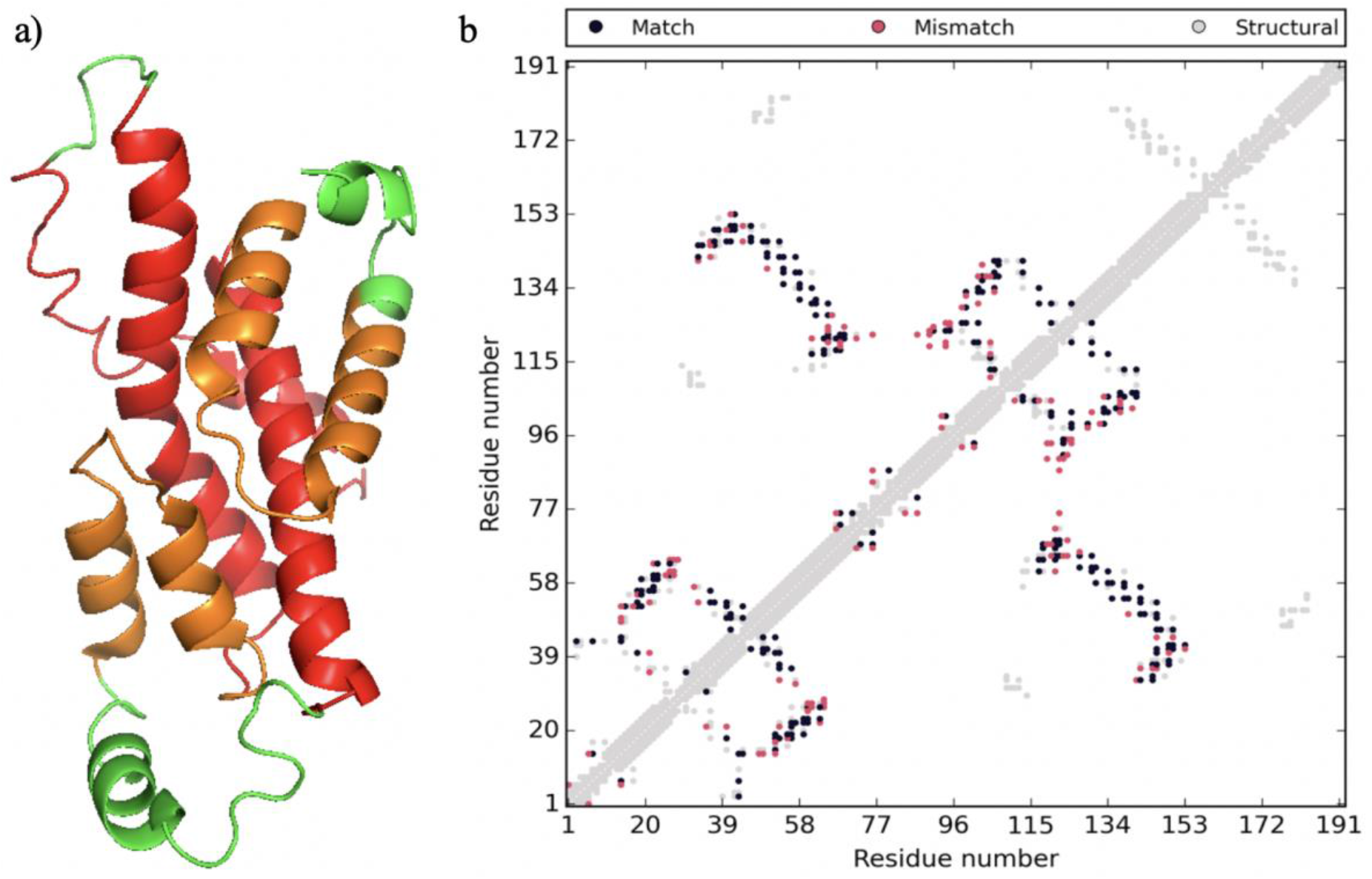
a) DMPfold model of Tmem41b with sequence conservation map (spectrum; blue (most conserved) to red (least conserved)) b) Comparison of model contact map with predicted contact map.

Again, the two halves of the model could be aligned with a TMscore of 0.5, indicating that the two fragments have the same fold. The duplicate fold is inserted into the membrane with opposing topology therefore resulting in the protein displaying an inverted repeat structure with 2-fold rotational symmetry about the membrane (Fig. 14).

**Fig 14.**
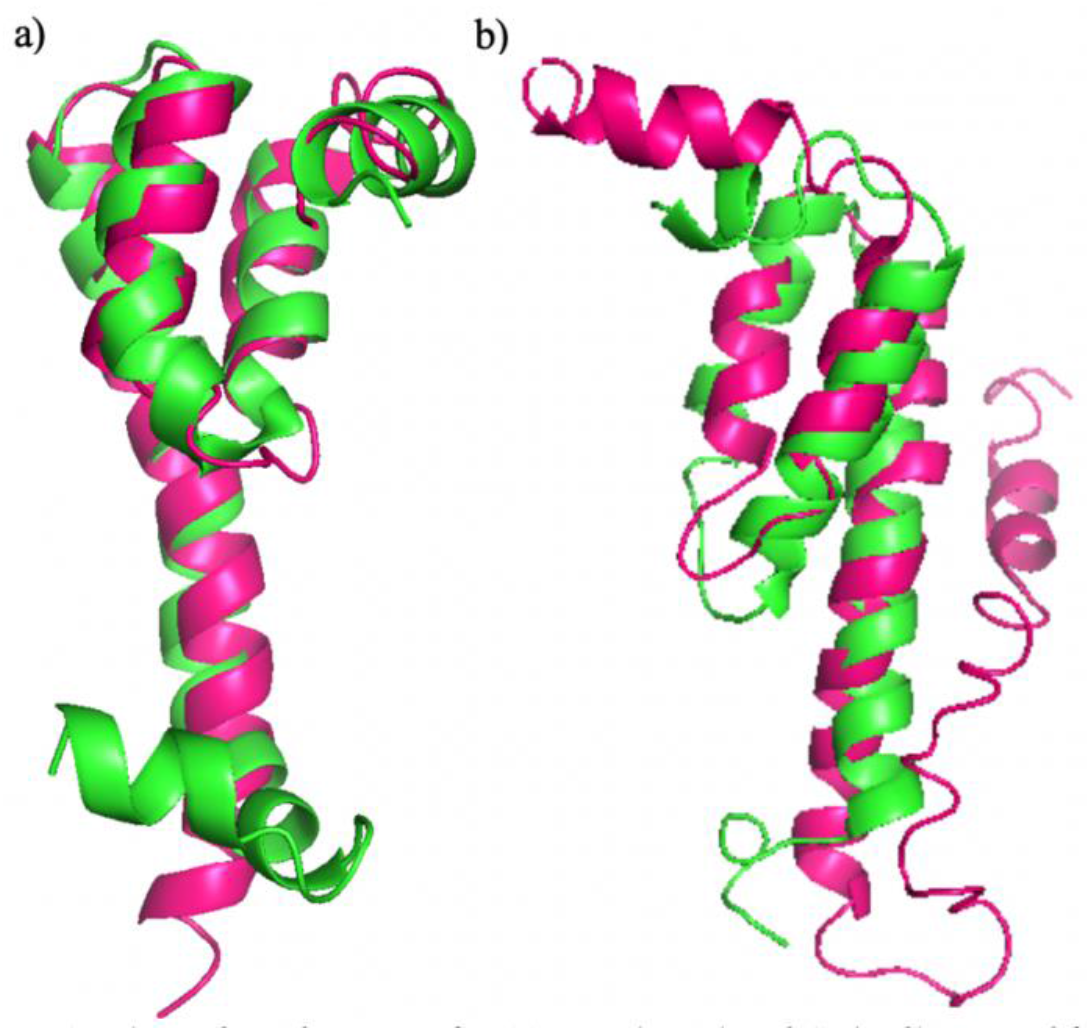
a) TMalign alignment of Mt2055 N- (green) and C- (pink) terminal fragments TMscore of 0.66 (normalised by C-terminal chain). b) TMalign alignment of Tmem41b N- (green) and C- (pink) terminal halves TMscore of 0.5 (normalised by N-terminal chain).

### Modelling confirms the presence of highly conserved proline residues at the turning point of the putative re-entrant helices

Assessing the conservation across the sequence of PF09335/PF06695 homologues, ConSurf highlights regions of strong conservation (Fig. 15). The strongly conserved regions are located in R1b/R2b & the mid-points of TM1/TM2 and come together in three-dimensional space. A similar distribution is witnessed when mapping conservation on to the pore region of the Cl^−^/H^+^ antiporter 3org (Fig. 15). In an effort to locate the presence of any functional residues the ConSurf data at the regions of highest conservation were examined. As expected from the sequence analysis above, the conservation data identified the presence of proline residues at the ‘turning point’ of all putative re-entrant loops in the MSAs. Thus, the conserved proline identified above is suggested by the models to have a structural role providing the tight turn required for the approximate 20° (Fig. 16) angle making up the re-entrant loop. Interestingly, there are examples where the presence of proline in a re-entrant loop allows it to act as a pivot, enabling the two segments of the loop to switch between states through a conformational change ^70,71^. Importantly, analysing the sequence of all re-entrant loops of the top cluster (comprising members of the Tmem41b family and the transporters; see above) revealed that they all contain a proline at the turning-point.

**Fig 15.**
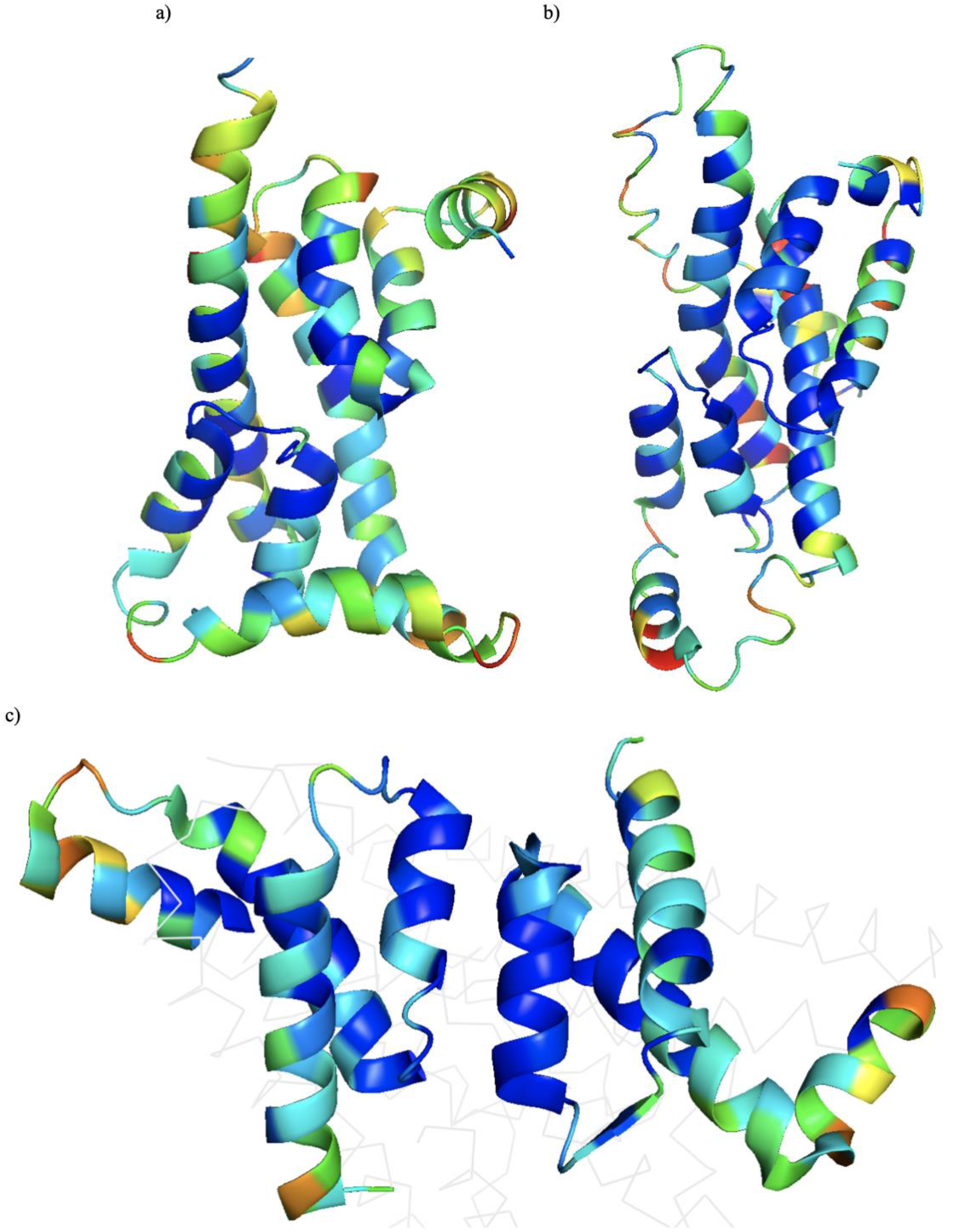
a) DMPfold model of Mt2055 with conservation map (spectrum; blue (most conserved) to red (least conserved)) b) DMPfold model of Tmem41b with conservation map (spectrum; blue (most conserved) to red (least conserved)) c) Crystal structure of 3org pore with conservation map (spectrum; blue (most conserved) to red (least conserved))

**Fig 16.**
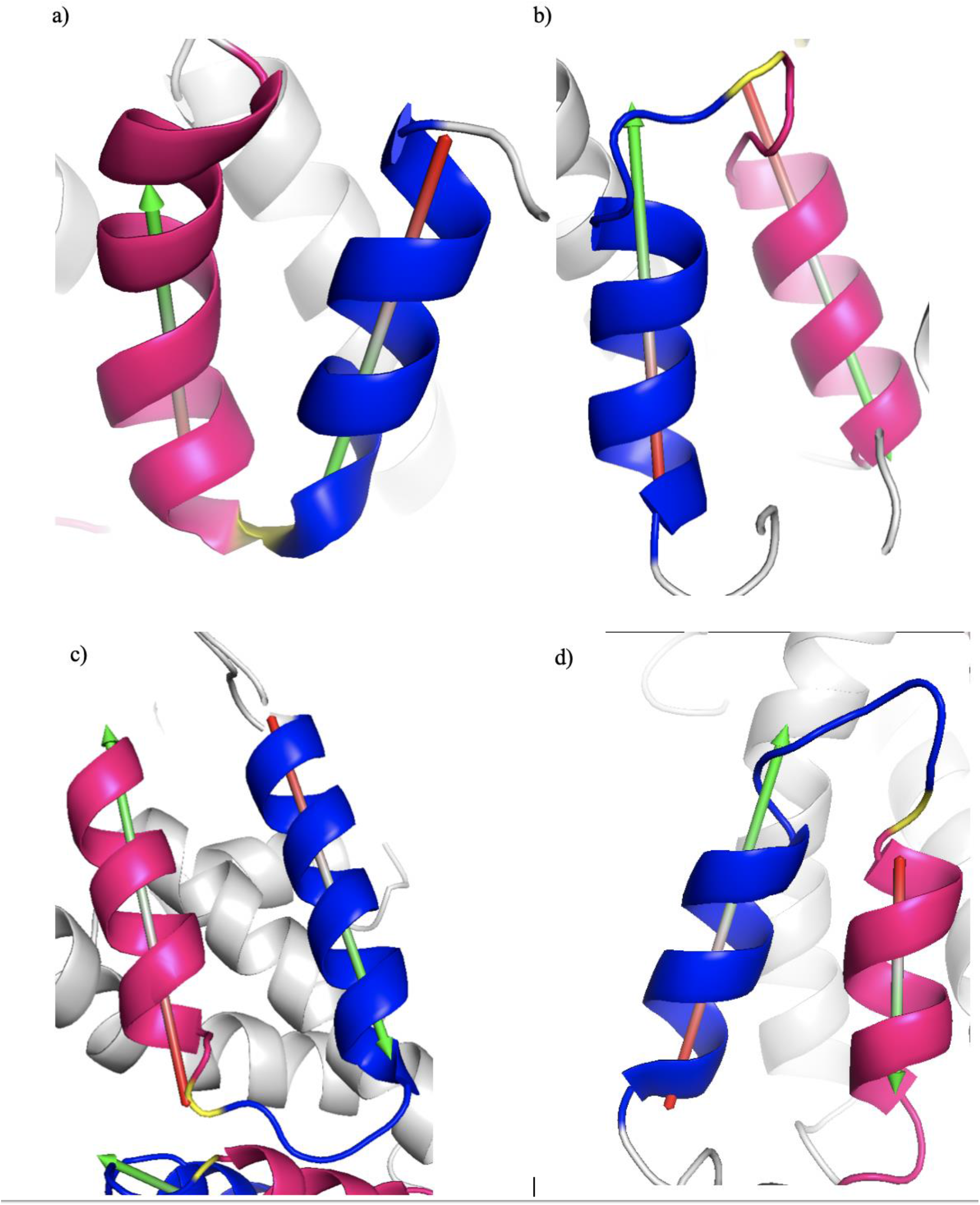
Tight re-entrant turn of around 160° a) Mt2055 R1 b) Mt2055 R2 c) Tmem41b R1 d) Tmem41b R2. Blue is C-terminal side of re-entrant loop, pink is N-terminal side of re-entrant loop, yellow is position of the proline. It can be seen that the proline residue is slightly off set from the turning point in some re-entrant loops possibly as a result of inaccuracy in the modelling.

## Conclusions

This study successfully demonstrates how covariance prediction data have multiple roles in modern structural bioinformatics: not just acting as restraints for model making and serving for validation of the final models, but also predicting domain boundaries and revealing the presence of cryptic internal repeats not evident by sequence analysis. This work highlighted a predicted contact map feature corresponding to packing of a re-entrant loop against a proceeding helix. Notably, structural alignment between our models and experimental structures of the region did not produce significant hits although the structural correspondence is clear from the covariance analysis.

Sequence, co-variance and predicted structure analysis shows that the Pfam PF09335 and the Pfam PF06695 domains are related. These domains contain a structural core composed of a pseudo-inverse repeat of an amphipathic helix, a re-entrant loop and a TM helix. All PF09335 homologues contain this central core with additional TM-helices flanking both sides of this core, in contrast to all homologues of PF06695, which are only made up of the amphipathic helix, a re-entrant loop and a TM helix core. The re-entrant loops studied here possess hydrophilic regions on the C-terminal side of these loops indicating that these structures are lining a membrane pore.

Querying the models against the PDB using Dali does not yield any significant hits. However, analysis of the prediction data reveals two features of Tmem41b and its homologues that independently suggest that they are secondary transporters: both an inverted repeat architecture and the presence of a re-entrant loop are independently and strongly associated with transporter function. Additionally, the fact Tmem41b and its homologues show structural similarities with H^+^ antiporters indicates that Tmem41b and its homologues may also couple substrate transport with an opposing H^+^ current.

Further work needs to be carried out, however, to determine what the substrate is for these putative transporters and confirm the mechanism of transport. The *ab initio* models do show highly conserved residues coming together in the region that would be buried in the membrane forming a substrate chamber consistent with the transport of a specific substrate.

## Supplementary Figures

**Supplementary Figure 1.**
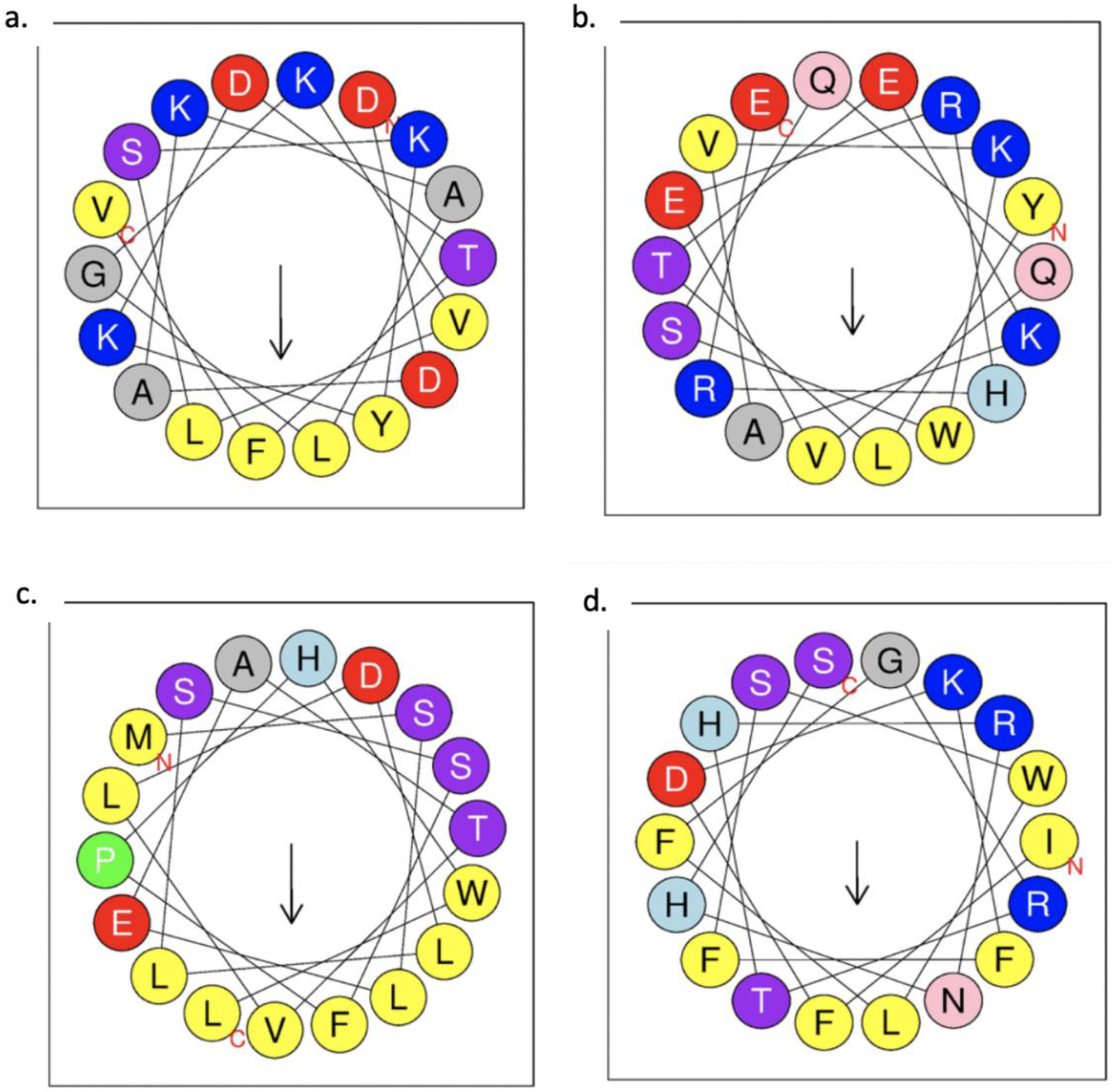
Helical wheel diagrams for putative amphipathic loops. Arrow representing direction hydrophobic moment with the size relative to magnitude of the moment; a. Tmem41b amphipathic loop 1; b. Tmem41b amphipathic loop 2; c. Mt2055 amphipathic loop 1; d. Mt2055 amphipathic loop 2

**Supplementary Figure 2.**
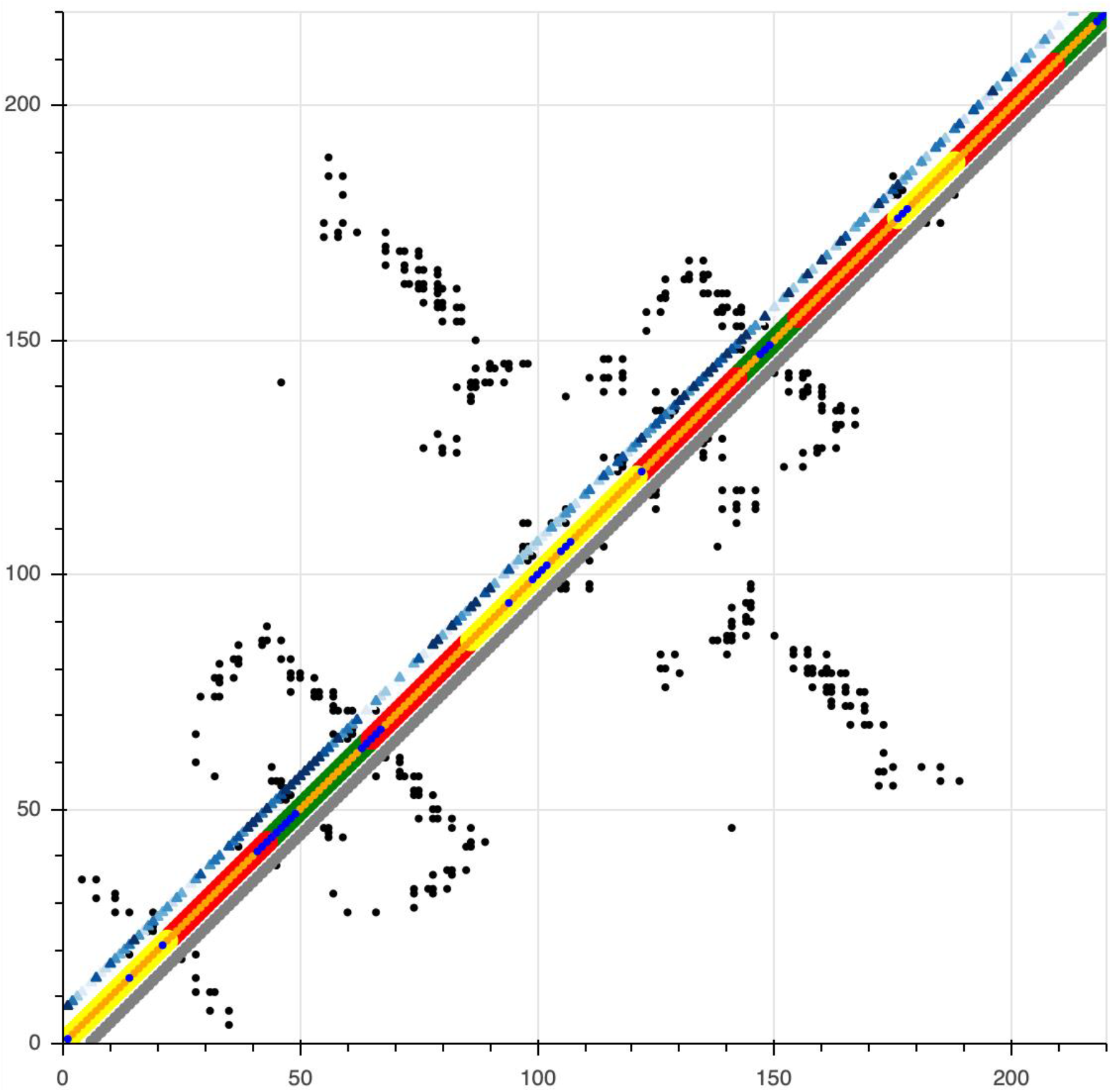
Yqja predicted contact map constructed using DeepMetaPSICOV with additional information overlaid on the diagonal. The outer diagonals show the TOPCONS membrane prediction (red regions being predicted TM helices, green; inside cell, yellow; outside). The thin central diagonal is the secondary structure prediction (orange, helix; blue, coil). Additionally, the upper diagonal contains a blue spectrum which indicates levels of sequence conservation from ConSurf (Ashkenazy et al., 2016) (the darker the blue the higher the level of conservation). Also, the grey (ordered) and lime green (disordered) lower diagonal utilises disorder predictions from IUPRED2a.

**Supplementary Figure 3.**
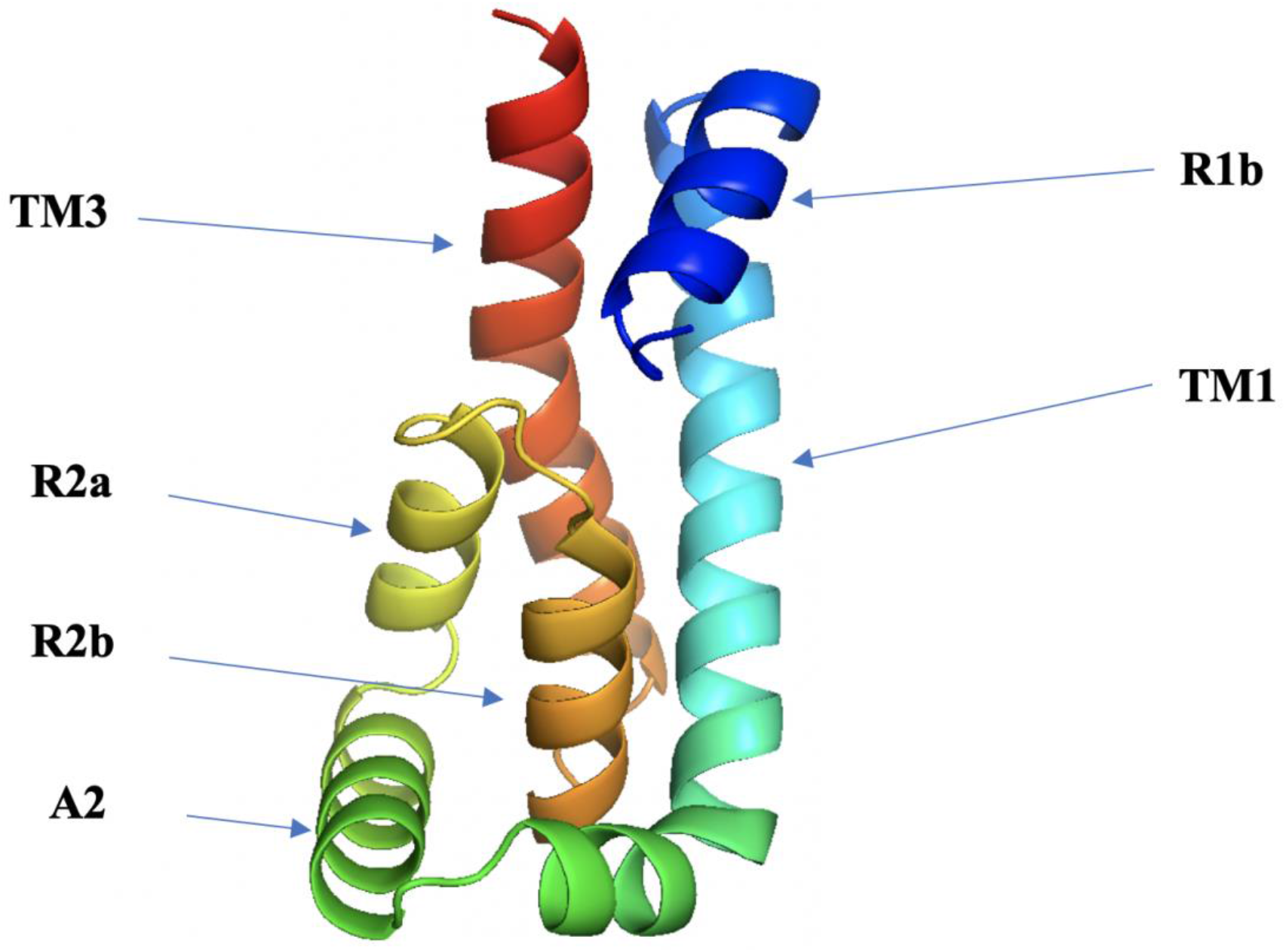
Automated DMPfold generated model for PF09335. The first amphipathic helix (A1) and the N-terminal half of the first re-entrant (R1a) is absent.

**Supplementary Figure 4.**
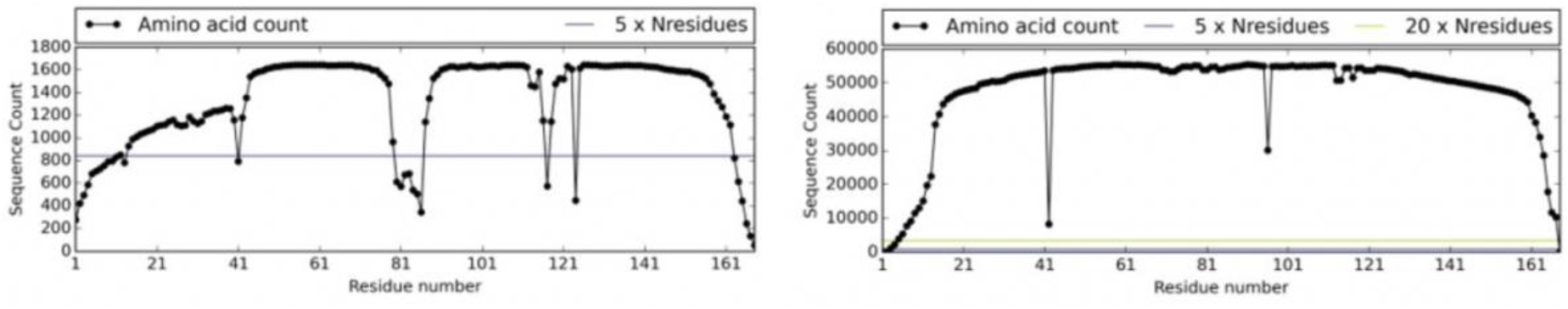
Sequence coverage for MSA of Mt2055 using UniRef90 (left) and sequence coverage for MSA of Mt2055 using metagenomic data (right).

**Supplementary Figure 5.**
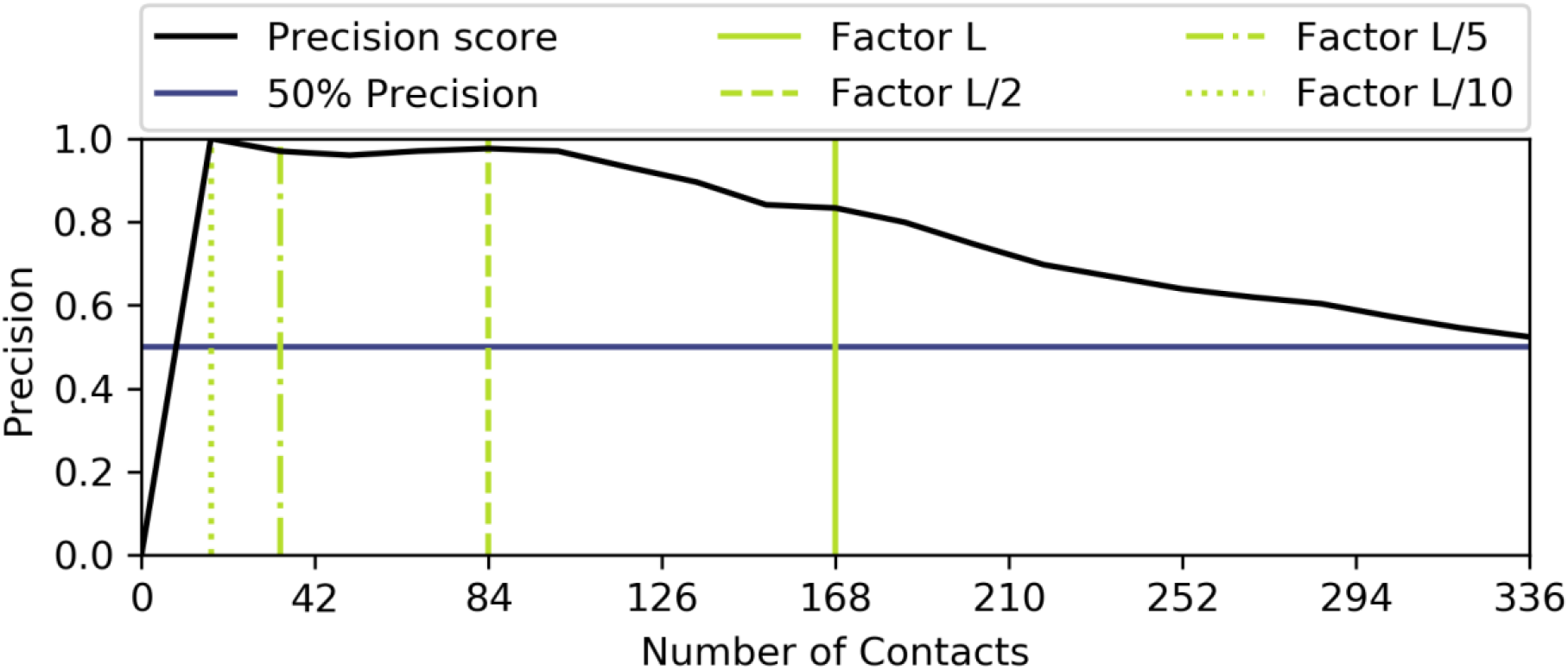
Precision scores for the Mt2055 ab initio model quantifying number of model contacts that have been satisfied by predicted contacts.

**Supplementary Figure 6.**
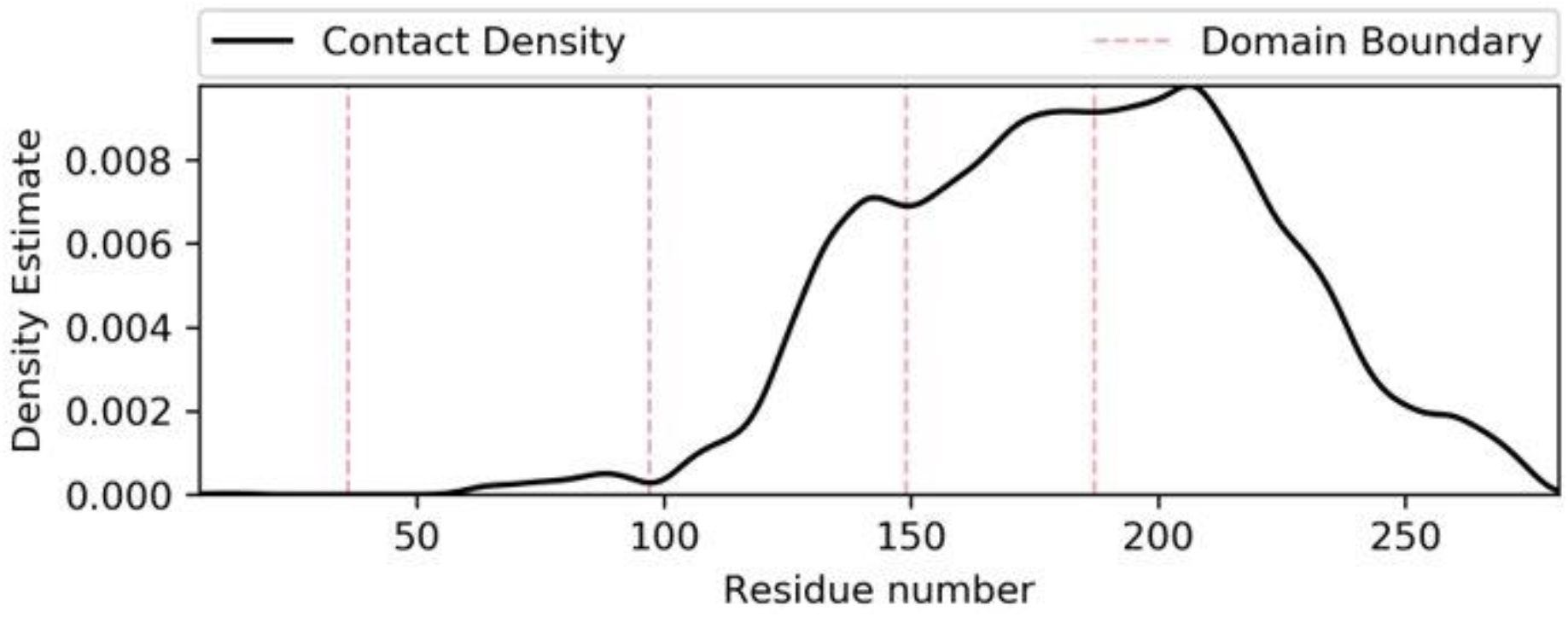
Predicted domain boundaries for Tmem41b based on predicted contact density. Disorder prediction data suggests that the first 40 residues is disordered. TM helix prediction data indicate the presence of a TM helix from positions 50-70 with the contact density profile showing that this region is not intimately packed with the rest of the protein.

**Supplementary Figure 7.**
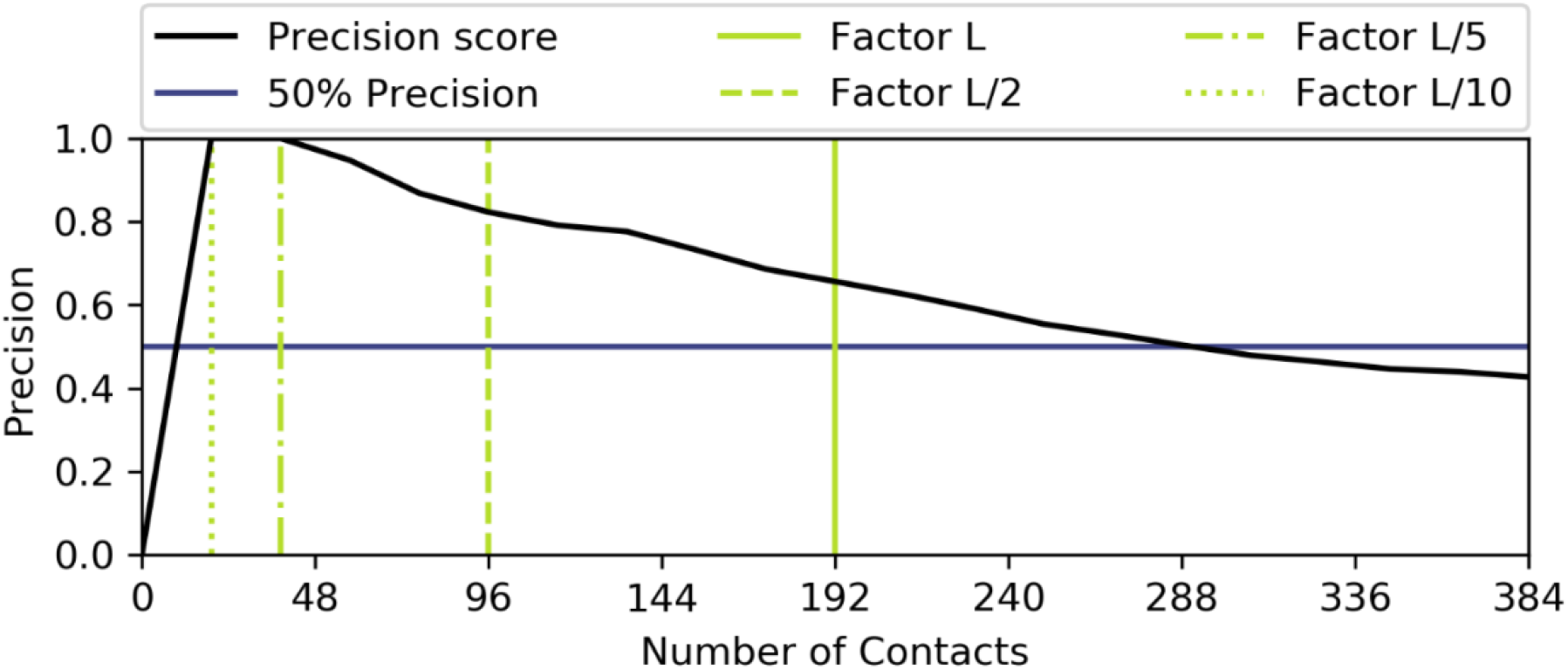
Precision scores for the truncated Tmem41b ab ignition model quantifying number of model contacts that have been satisfied by predicted contacts.

